# SARS-CoV-2 Defective Viral Genomes from Distinct Genomic Regions Drive Divergent Interferon Responses

**DOI:** 10.64898/2026.03.19.712870

**Authors:** Justin W. Brennan, Simone Spandau, Xingjian Wang, Haley Aull, Sarah Connor, Gloria Pryhuber, Thomas J. Mariani, Ruth Serra-Moreno, Yan Sun

**Author notes:** **Correspondence:**, 601 Elmwood Avenue Rochester, NY 14642.

## Abstract

Defective viral genomes (DVGs) are naturally generated during genomic replication of many RNA viruses. When produced early in infection or supplemented at the onset of infection, DVGs can attenuate viral pathogenesis by stimulating IFN responses and antagonizing wild type (WT) virus replication, highlighting their potential as antiviral therapeutics. However, during natural infection DVGs can exert both antiviral and proviral effects depending on their generation kinetics, species, and abundance, underscoring the need to better understand their roles in viral pathogenesis. Coronaviruses (CoVs) remain a major global health threat and ubiquitously generate DVGs, yet DVGs’ roles during CoV infection are largely unknown. Using SARS-CoV-2 as a model, we previously identified DVGs in vitro and in patient samples and discovered two major genomic hotspots (A and B) for their generation. Here, we first showed that overall DVG abundance tended to positively correlate with COVID-19 severity, with approximately 40% of DVGs originating from a specific genomic region designated hotspot B. Analysis of a publicly available single-cell RNA-seq datasets revealed that DVGs from hotspot B, but not hotspot A, were associated with elevated IFN responses, suggesting that DVGs derived from different genomic regions vary in their ability to stimulate innate immunity. To test this directly, we constructed two representative DVGs corresponding to hotspots A and B. Both DVGs suppressed the replication of co-infecting WT virus; however, only DVG-B induced robust IFN responses, exceeding those triggered by WT virus alone. This was further confirmed in human precision-cut lung slices. Mechanistically, DVG-B–derived dsRNA exhibited a distinct subcellular distribution compared with WT virus. Complementation with the nucleocapsid (N) partially restored dsRNA organization but did not alter the IFN response. Together, our findings demonstrate that DVGs arising from distinct genomic hotspots differentially regulate IFN responses, potentially contributing to varied pathogenic outcomes during SARS-CoV-2 infection.

**Author summary:** Defective viral genomes (DVGs) are naturally produced during RNA virus infection and can suppress viral pathogenesis by stimulating innate immune responses. However, their roles in coronavirus infection, particularly SARS-CoV-2, remain poorly understood. This study investigated the species-specific function of DVGs generated during SARS-CoV-2 infection and their impact on disease outcomes. Cohort analysis revealed that overall DVG abundance tended to positively correlate with COVID-19 severity, with approximately 40% of DVGs originating from a specific genomic region designated hotspot B. Single-cell RNA sequencing showed that DVGs from hotspot B, but not hotspot A, were associated with elevated IFN responses, suggesting that DVGs from different genomic regions vary in their immunostimulatory capacity. To directly test this, we constructed representative DVGs from both hotspots. While both suppressed wild-type virus replication, only DVG-B induced robust IFN responses both *in vitro* and *ex vivo*. Mechanistically, DVG-B produced dsRNA with distinct subcellular distribution compared to wild-type virus. Interestingly, complementation with the viral nucleocapsid protein partially restored dsRNA organization but did not alter IFN responses. These findings demonstrate that SARS-CoV-2 DVGs arising from different genomic hotspots differentially regulate innate immunity, potentially contributing to varied pathogenic outcomes during infection.

## Introduction

Defective viral genomes (DVGs) are naturally generated by most RNA viruses and play important roles in viral pathogenesis[1–14]. When arising early during infection, DVGs can reduce viral pathogenicity by inducing robust type I and type III interferon (IFN) responses and compete for critical viral proteins required for genomic replication and packaging to effectively suppress wild-type (WT) virus replication[15]. For this reason, DVGs are being explored as a novel approach for antiviral therapeutics. However, when DVGs emerge later during infection (during and after the peak viral load), their presence is associated with enhanced inflammation, higher viral titers, and worse symptom severity in patients[15]. Additionally, for some viruses, DVGs have been shown to facilitate viral persistence[16, 17]. Together, these findings suggest that DVGs can have both antiviral and proviral effects depending on their kinetics of generation, species, and abundance. Further investigations into DVG function and their impact on viral pathogenesis are needed to harness their potential to mitigate virus-induced diseases.

Severe acute respiratory syndrome coronavirus 2 (SARS-CoV-2) is the causative agent of the coronavirus disease 2019 (COVID-19) pandemic, which resulted in extensive morbidity and mortality worldwide. Although the acute phase of the pandemic has subsided, SARS-CoV-2 continues to circulate globally and remains a significant public health concern. Continued investigation into the viral factors contributing to SARS-CoV-2 pathogenicity and how the virus interacts with host immune responses are therefore required. SARS-CoV-2 belongs to the *Betacoronavirus* genus and contains a positive-sense ∼30 kb RNA genome[18]. Roughly two-thirds of the genome encodes 16 nonstructural proteins that form the viral polymerase complex responsible for viral replication and transcription. The remaining one-third encodes structural and accessory proteins. Among these, the nucleocapsid (N) protein is essential for ribonucleocapsid formation and virus particle assembly[19]. Compared to influenza virus, another pandemic-causing virus, SARS-CoV-2 induces relatively weak interferon (IFN) responses *in vitro*[20]. During infection, SARS-CoV-2 generates double-stranded RNA (dsRNA) as a replicative intermediate of genomic replication and subgenomic mRNA transcription. These dsRNAs can be sensed by RIG-I–like receptors (RLRs), leading to the induction of type I and type III IFNs through mitochondrial antiviral-signaling protein (MAVS) activation[21, 22]. However, viral dsRNAs are typically confined within virally generated double-membrane vesicles (DMVs), limiting their exposure to host dsRNA sensors. In addition, SARS-CoV-2 encodes multiple IFN antagonists, such as nsp1, N, and several accessory proteins, which further block IFN production and signaling pathways[22]. Despite these immune evasion strategies, elevated IFN responses have been observed in some COVID-19 patients, and early robust IFN responses are associated with improved clinical outcomes[23]. These observations highlight the importance of understanding the viral and host factors that regulate IFN induction during SARS-CoV-2 infection.

DVGs are ubiquitously generated during SARS-CoV-2 infections *in vitro* and in patients[17, 24–28]. SARS-CoV-2 primarily generates deletion type DVGs that harbor internal deletions ablating essential genes required for virus replication or assembly. We previously demonstrated that the abundance of SARS-CoV-2 DVGs is positively correlated with type I IFN responses. In addition, symptomatic COVID-19 patients were found to harbor higher levels of DVGs than asymptomatic individuals, suggesting that DVGs may influence IFN responses and contribute to differences in COVID-19 outcomes. Importantly, we previously identified three genomic hotspots for DVG generation, designated A, B, and C. Among these, hotspots A and B represent the two major sites contributing to the SARS-CoV-2 DVG population. Hotspot A is primarily observed *in vitro* and contains a large deletion ablating nearly the entire viral genome, leaving the 5’ and 3’ untranslated regions (UTRs) and a small, truncated fragment of nsp1. As DVGs in hotspot A lack nonstructural proteins, they cannot independently replicate in the absence of WT SARS-CoV-2. In contrast, hotspot B harbors a 2–3 kb deletion region near the 3′ UTR, eliminating ORF7a/7b, ORF8, and N. Because N is essential for virus particle assembly and all 16 nonstructural proteins are intact in hotspot B DVGs, they are capable of independent genomic replication but deficient in virus particle assembly. Moreover, hotspot B is the most conserved among the three and is detected both *in vitro* and in samples from COVID-19 patients[17].

Here, we aimed to examine the functions of DVGs from two major hotspots A and B, specifically their ability to induce IFN and suppress WT virus replication. Using a publicly available cohort, we found a tendency that total DVG abundance positively correlated with COVID-19 disease severity. Consistently, approximately 40% of those DVGs were in hotspot B. We then utilized a single-cell (sc) RNA-seq dataset from *in vitro* infection and observed that DVGs in hotspot B, but not hotspot A, were associated with elevated IFN responses, suggesting DVGs from different genomic regions differ in their ability to stimulate IFN. To directly test this, we constructed two representative DVGs using the outermost recombination boundaries in hotspot A and B to maximally encompass the deletions in these two DVG populations. Functional analysis indicated that both DVGs suppressed the replication of a co-infected WT virus, while only DVG-B induced robust IFN responses, significantly greater than the level induced by the WT virus alone. This observation was further verified in human precision-cut lung slices (PCLS) from 4 different adult donors. Mechanistically, DVG-B exhibited a distinct subcellular distribution of dsRNA compared to the WT virus. Interesting, complementation of N partially restored DVG-B’s dsRNA pattern to that of WT. This, however, did not correspondingly alter the IFN response induced during DVG-B infection. These data indicated that N was critical in dsRNA/DMV organization, but the absence of N was not the primary driver of the robust IFN induction triggered by DVG-B. Taken together, our data indicate that DVG-A and DVG-B differ in their IFN induction capacities, potentially driving distinct pathogenesis outcomes during SARS-CoV-2 infection.

## Results

### SARS-CoV-2 DVGs tended to positively correlate with COVID-19 disease severity

We first sought to examine whether DVGs impact infection outcomes in COVID-19 patients. We obtained one published scRNA-seq dataset from nasal swabs collected from a COVID-19 patient cohort with known severity scores and age information (SCP1289[29]). Using our previously published pipeline[17], we observed ∼35% (13/37) of COVID-19 patients were positive for DVGs (Fig. 1A). No DVGs were identified in COVID-19 negative patients. DVG+ patients had significantly higher viral read counts than DVG-patients and showed a trend toward higher severity scores. No age differences between the DVG+ and DVG-groups was observed (Fig. 1B). Because age is an important factor impacting COVID-19 severity, we divided the COVID-19 patients into two age groups: younger (age<65) and elderly (age>65) and compared their DVG amount, viral load, and disease severity. We observed that, on average, elderly patients had significantly more total DVGs than younger patients associated with higher severity scores (Fig. 1C). Correlation analysis of 37 COVID-19 patients revealed a weak positive correlation between DVG abundance and disease severity scores. This trend was more evident for DVG abundance compared to viral load (Fig. 1D). Previously, we identified two major hotspots where SARS-CoV-2 DVGs were produced, hotspot A and B, via bulk RNA-seq[17]. Hotspot A was observed primarily *in vitro*, whereas hotspot B was conserved among *in vitro* infections and autopsy lung tissues of COVID-19 patients. From this cohort study, we consistently observed that hotspot B was more prevalent than hotspot A across both age groups, accounting for ∼40% of the total DVG population (Fig. 1E). Collectively, our analyses indicate that despite a weak correlation between SARS-CoV-2 DVGs and disease severity, hotspot B represents a dominant site of DVG generation compared with other genomic regions, warranting further mechanistic investigation.

**Fig 1.**
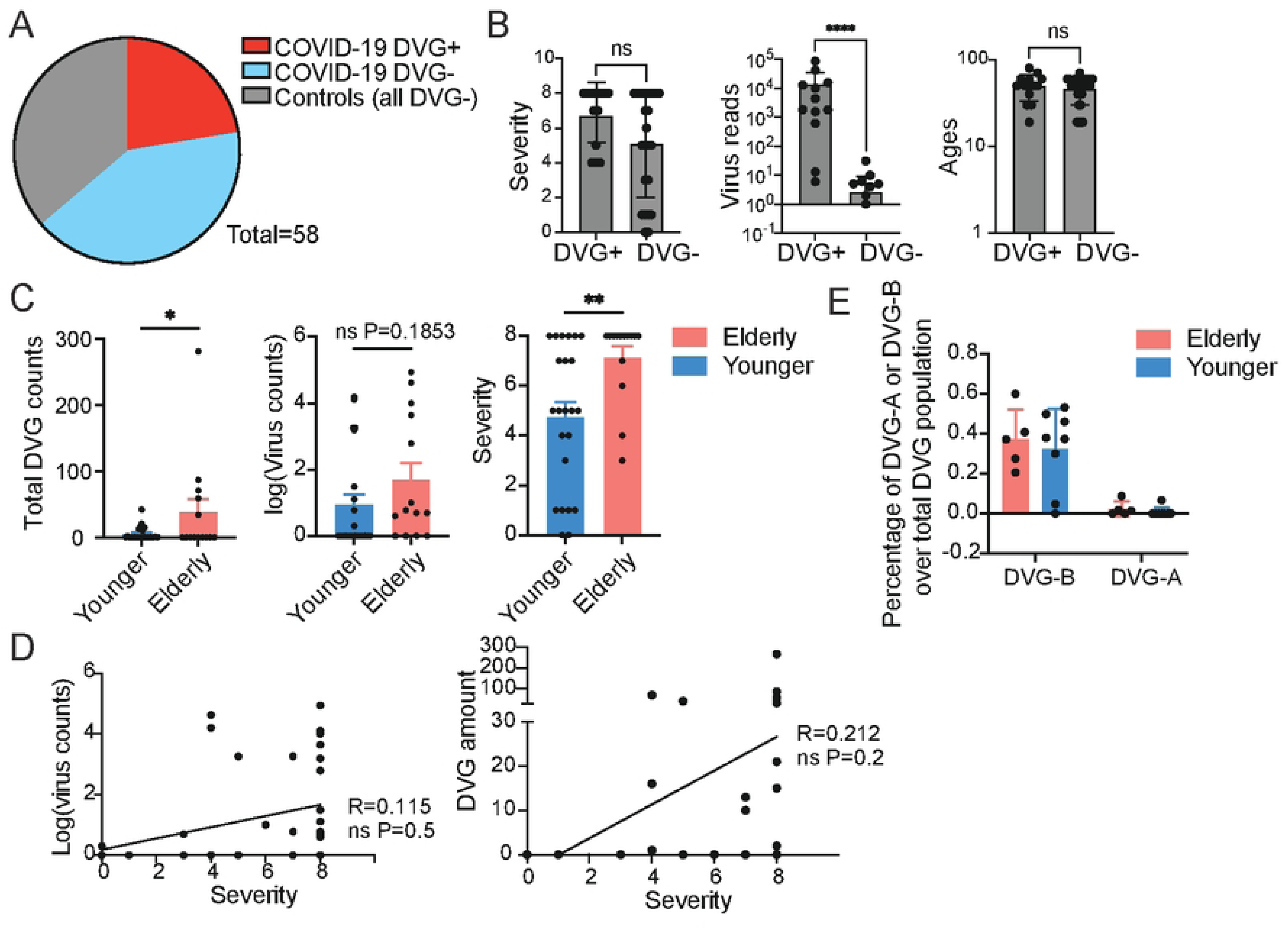
SARS-CoV-2 DVGs positively correlated with COVID-19 disease severity. **(A)** The pie chart shows the proportions of DVG- and DVG+ COVID-19 patients, along with control patients, in the cohort. **(B)** Bar graphs shows the comparisons of disease severity score, virus reads, and ages between DVG+ and DVG- COVID-19 patients. **(C)** Bar graphs compared the total DVG read counts, viral counts, and severity score between younger (<65 yrs old) and elderly (>65 yrs old) COVID-19 patients. * P<0.05, ** P<0.01, **** P<0.0001 by Mann-Whitney test. mean±SD (**D**) showing the correlation between total viral reads counts or DVG read counts and disease severity in COVID-19 patients. * P<0.05 by Pearson r correlation. **(E)** The bar graph displayed the percentages of DVGs in hotspots A and B among DVG+ COVID-19 patients from two different age groups.

### DVGs clustered in hotspot B, but not hotspot A, were associated with the induction of type I and type III IFNs

As a population, SARS-CoV-2 DVGs have been associated to enhance type I and type III IFN production during *in vitro* infection[17]. To examine whether DVGs in different hotspots equally contribute to IFN responses, we analyzed one publicly available scRNA-seq dataset using primary adult human bronchial epithelial cells infected at MOI of 0.01 (GSE166766[30]). From this dataset we previously found that DVGs were majorly detected at days 2 and 3 post infection[17]. Focusing on 2 days post-infection (2 dpi), we plotted DVG+ cells according to their DVG junction positions (E’ and V’ in Fig. 2A) and examined their association with single-cell expression levels of IFNB1, IFNL1, and Mx1. Surprisingly, we found that IFN high-expressing cells (red dots) were clustered in DVG hotspot B but completely absent in hotspot A (Fig. 2B). The same trend was observed in three cell types containing the greatest amount of DVGs (Fig. 2C). Although there were more cells positive for hotspot B than hotspot A, the average viral counts and DVG counts per cell remain the same between the two (Fig. 2D). A similar trend was observed for samples at 3dpi (Fig. S1). Together, these results strongly suggest that DVGs in hotspot B, but not hotspot A, contribute to the induction of IFN responses during infection.

**Fig 2.**
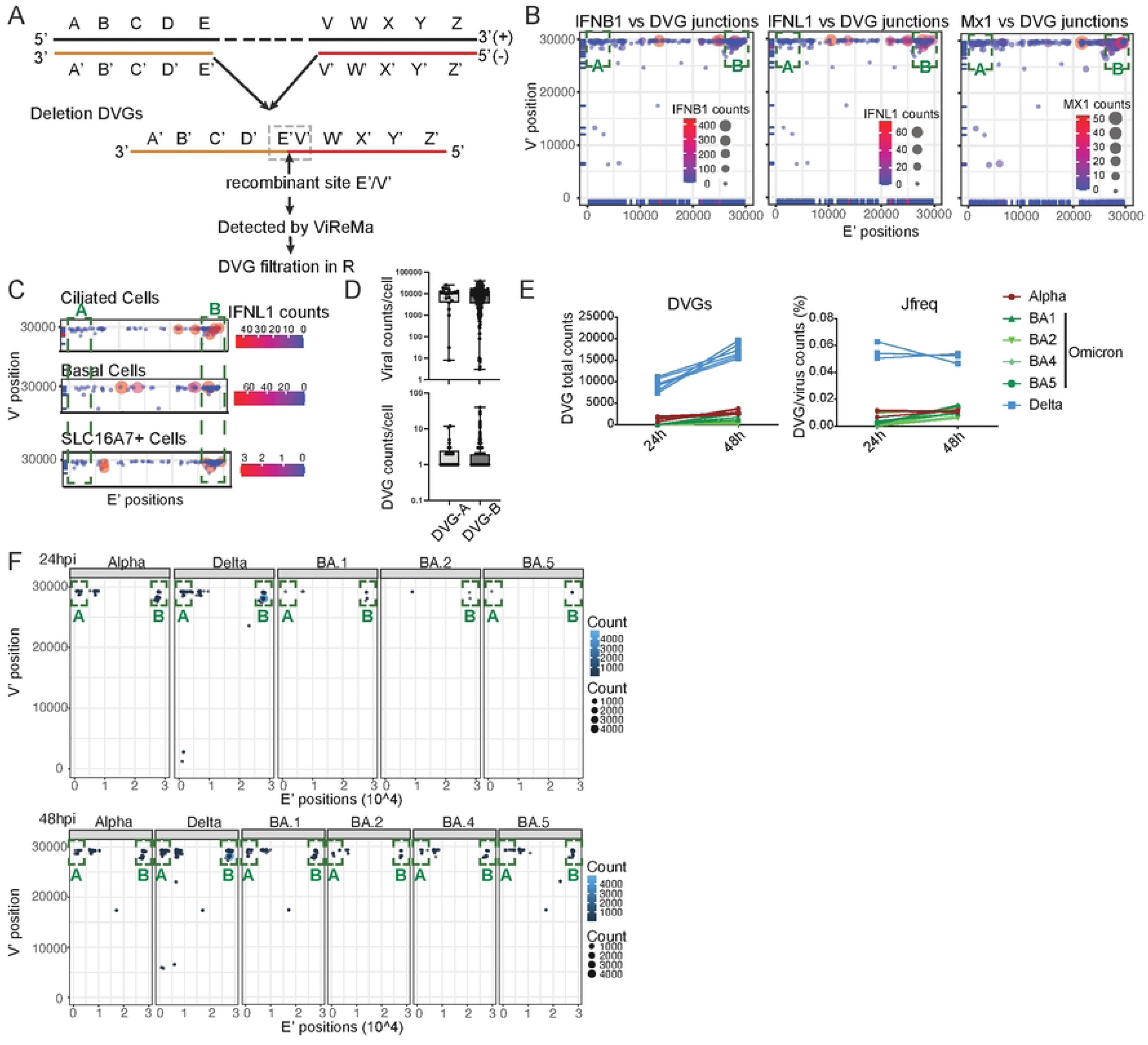
DVGs clustered in hotspot A and B showed different IFN induction ability via single-cell RNA-seq. **(A)** Schematic representation of DVG generation from the positive sense viral genome and the general principle of ViReMa identification of deletion DVGs. The V’ site represents the break point and the E’ site represents the rejoin point of the viral polymerase in the formation of DVGs. The gray dashed box marks the recombinant site that distinguishes DVGs from full length viral genomes, which are identified by ViReMa, and further filtered using R script published previously. **(B)** Gene expression of IFNB1, IFNL1, and Mx1 were correlated with DVG positions at a single cell level at 2 dpi in NHBE cells (GSE166766). Each dot represented individual DVG+ cell. Both the color and size of dots were based on the expression level of indicated genes. Green bracket indicated the previously identified hotspot A and B. **(C)** Within three major cell types positive for DVGs, gene expression of IFNL1 (2dpi) were correlated with DVG positions at the single cell level similarly as shown in (B). **(D)** At 2dpi, the viral reads and DVG junction reads in cells positive for DVGs at hotspot A (DVG-A) were compared with those in cells positive for DVGs at hotspot B (DVG-B). DVG junction hotspots among different SARS-CoV-2 variants were then analyzed using the publicly available dataset GSE213759. **(E)** The respective total DVG junction read counts, and J_freq_ percentages were graphed for different variants of concerns: Alpha, Delta and several Omicrons at 24h and 48h post infections in Calu3 cells. **(F)** showing break point (V’) and rejoin point (E’) distributions for DVGs from (E) with upper panel for 24h and lower pannel for 48h. Circle size and color intensity indicated the DVG counts. Green bracket indicated the previously identified hotspot A and B.

### DVG junction hotspots were conserved among different SARS-CoV-2 variants

Because we observed different IFN stimulatory abilities by DVGs from hotspot A and hotspot B, we asked whether these two hotspots are conserved amongst different SARS-CoV-2 variants. To do this, we obtained another publicly available NGS dataset (GSE213759), wherein Calu3 cells were infected with different variants of concern (VOC) at the same MOI[31]. We specifically analyzed DVG amounts and their genomic locations at 24h and 48h post infection. We detected greater total DVG amount and Jreq (DVG reads normalized by total viral reads) in the Delta variant compared with other variants (Fig. 2E). This higher level may reflect a more efficient production and/or enrichment of DVGs in the Delta variant virus stocks during passaging and preparation. Importantly, all variants showed DVG hotspot A and B at 24- or 48-hpi (Fig. 2F), indicating that these two hotspots were conserved among different variants during *in vitro* infection.

### Synthetic DVG-A and DVG-B particles attenuated WT virus replication, but only DVG-B stimulated robust IFN responses

To directly examine the species-specific function of SARS-CoV-2 DVGs during infection, we synthesized SARS-CoV-2 virus-like particles (VLPs) specifically packaging a representative DVG from hotspot A, DVG-A, and hotspot B, DVG-B, respectively. Based on the outermost range of hotspot A recombination junctions, DVG-A harbors a deletion ablating genomic positions 700-29,674. Therefore, we constructed a DVG-A plasmid by replacing this deletion with the mKate2 gene and an identified virus packaging signal located in nsp15 and nsp16 (PS9)[32] (Fig. 3A, right scheme). We then co-transfected the DVG-A plasmid with plasmids encoding the four SARS-CoV-2 essential structural proteins (E, M, S, and N) in 293T cells as previously reported[32], followed by purification using a 30% sucrose cushion. DVG-A titers were determined by RT-qPCR (Fig. 3A, left). The representative DVG-B harbors a deletion spanning genomic positions 27,700-29,674. This deletion region encodes the orf7a/b, orf8 and N genes. Among these genes, only N is essential, required for virus particle assembly. This deletion was introduced by replacing these viral sequences with the eGFP gene (under the N TRS-B sequence) in the corresponding circular polymerase extension reaction (CPER) fragment and constructed the full-length DVG-B via CPER approach[33] (Fig. 2A, right scheme). We then co-transfected the CPER products with plasmids encoding the viral structural proteins in HEK293T cells. Supernatants were filtered, concentrated, and subsequently passaged three times in VeroE6 cells constitutively expressing the N protein (VeroE6:N), followed by sucrose purification and titered by GFP FFU/ml determination in VeroE6:N cells (Fig. 3A, left). To characterize DVG-A, we infected VeroAT cells and BSRT7 cells (which lack ACE2) with either the DVG-A stock (DVG-A+) or the supernatants from HEK293T cells transfected with the DVG-A encoding plasmid alone, without the essential structural proteins (DVG-A-). RT-qPCR was performed on infected cells at 24 hpi. DVG-A genomes were detected only in DVG-A+ infected VeroAT cells, indicating that DVG-A genomes were successfully packaged into SARS-CoV-2 like viral particles (Fig. 3B). To characterize DVG-B, VeroAT cells were infected with supernatants from HEK293T cells co-transfected with the DVG-B CPER products and various combinations of the four essential structural protein encoding plasmids. Expression of all four structural proteins yielded the highest percentage of GFP+ cells, followed by the condition lacking Spike. In contrast, transfection with E, M, and S but without N produced no GFP signal, given that DVG-B does not encode N and is therefore unable to assemble virus particles (Fig. 3C). To further confirm that DVG-B particles cannot produce infectious virions in cells without N expression, VeroAT cells were infected with DVG-B, and the resulting supernatants were then used to infect new VeroAT cells. As expected, no GFP+ cells were detected by flow cytometry (Fig. 3D, left). In contrast, serial passaging of DVG-B in VeroE6:N resulted in GFP+ cells (Fig. 3D, right). To assess the expression of other viral proteins encoded by DVG-B, A549-ACE2 cells were infected with WT virus (Hong Kong strain, MOI=1), DVG-B (MOI=1), or DVG-A (MOI=1). The spike protein was readily detected in DVG-B infected cells, but not DVG-A infected cells via immunofluorescent assay (IFA) and western blot (Fig. 3E and 3F). Spike protein levels in DVG-B–infected cells were markedly lower than in WT virus–infected cells (Fig. 3F).

**Fig 3.**
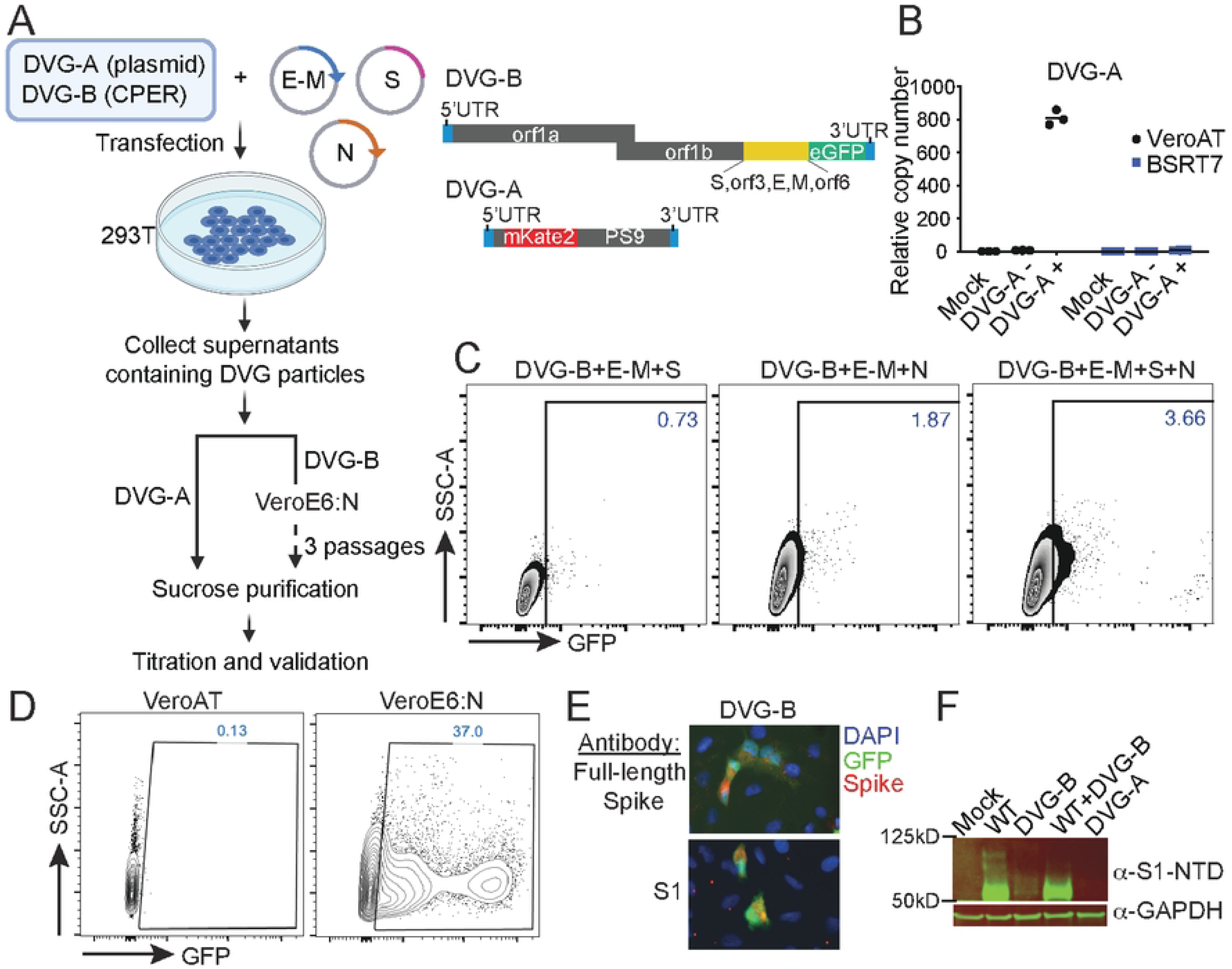
Generation and characterization of presentative DVG particles in hotspot A (DVG-A) and hotspot B (DVG-B). **(A)** Schematic representation of structures and production of DVG-A and DVG-B. Briefly, For DVG-B, genomic positions 27,700-29,674 were replaced with eGFP and rejoined to the remaining viral genome using CPER. For DVG-A, genomic positions 700-29,674 were replaced with the mKate2 reporter and PS9 and cloned into a plasmid under the CMV promoter. The DVG-B CPER product or DVG-A plasmid was co-transfected with four essential structural proteins, E, M, S, and N encoding plasmids into 293T cells. Supernatants were collected. For DVG-A particles, final stock was purified and concentrated by 30% sucrose cushion. DVG-B particles were further passaged in VeroE6:N cells for three passages to increase titer, followed by sucrose purification. **(B)** VeroAT and BSRT7 cells (no ACE2, negative control) were infected with mock, supernatants from 293T cells transfected with the DVG-A plasmid alone, without the essential structural proteins (DVG-A-), or the DVG-A stock (DVG-A+). RNAs were extracted at 24 hpi followed by RT-qPCR targeting specific sequences in DVG-A RNAs. **(C)** 293T cells were transfected with DVG-B CPER products and different combinations of E, M, S, and N. Supernatants were collected and used to infect VeroAT cells. Representative flow plots show GFP+% VeroAT cells infected with indicated supernatants. **(D)** DVG-B was used to infect VeroAT or VeroE6:N cells. Subsequent supernatants were passaged in VeroAT or VeroE6:N cells. Infected cells were analyzed by flow cytometry to determine the percentage of GFP+ cells at 3dpi. **(E)** A549-ACE2 cells were infected with DVG-B (MOI=1), followed by IFA using two spike antibodies: one targeting the full-length spike (top) and one tarting the S1 region (bottom). **(F)** A549-ACE2 cells were infected with WT SARS-CoV-2 (HK, MOI=1), DVG-B (MOI=1), DVG-B+WT (ratio 1:1), and DVG-A (MOI=1). Cells were collected at 24hpi, followed by western blotting and blotting by Spike antibody targeting S1-NTD.

We next used characterized DVG-A and DVG-B (passage 3, P3) stocks to assess their direct effects on WT virus infection and IFN responses. A549-ACE2 cells were infected with WT virus (MOI = 1), DVG-A (MOI = 1), DVG-B (MOI = 1), WT + DVG-A (1:1), or WT + DVG-B (1:1). Cells were collected at 24 hpi for RT-qPCR analysis of IFN-related and viral gene expression. WT virus infection induced minimal IFN responses compared to mock infection. Despite reduced replication, DVG-B infection alone elicited significantly higher IFN responses than WT virus infection. When co-infected with WT virus, DVG-B induced IFN responses were further enhanced (Fig. 4A). IFNB1 ELISA of A549 supernatants confirmed the induction of IFNB1 by DVG-B alone at protein level (Fig. 4C). In contrast, DVG-A did not induce any IFN responses (Fig. 4B). TCID₅₀ assays showed that both DVG-A and DVG-B suppressed WT virus replication in a dose-dependent manner (Fig. 4D-F). The suppressive effects plateaued at DVG:WT ratios of 5:1 for both DVG-A and DVG-B (Fig. 4E–F).

**Fig 4.**
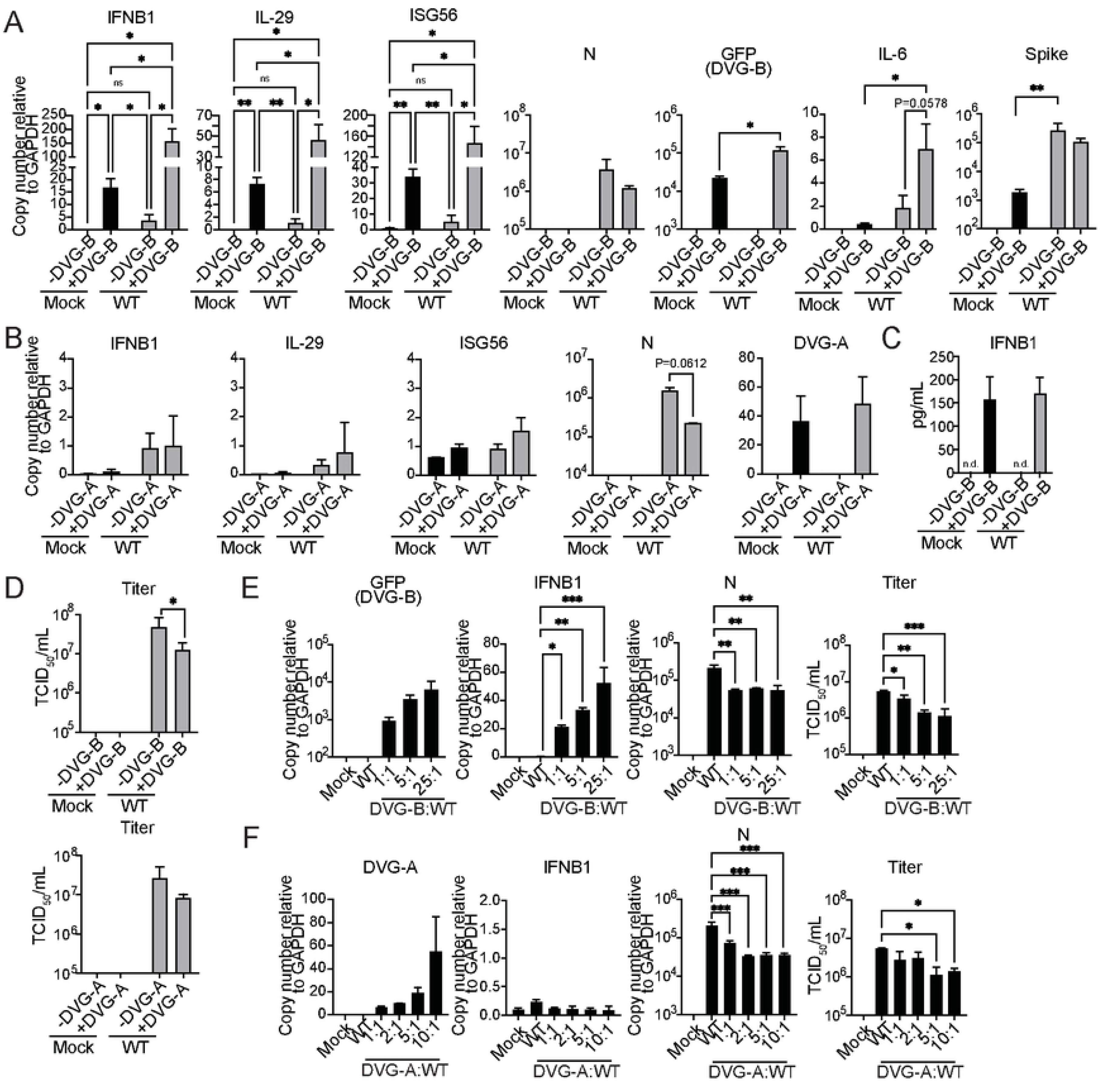
DVG-A and DVG-B containing virus-like particles antagonize WT SARS-CoV-2 infection. **(A-B)** A549-ACE2 cells were infected with SARS-CoV-2 (MOI 0.8), DVG-B (**A**, MOI 1), DVG-A (**B**, MOI 1) or co-infected with either DVG-B:SARS-CoV-2 (MOI 1: MOI 0.8) or DVG-A: SARS-CoV-2 (MOI 1: MOI 0.8) for 24 hours post-inoculation. Expression of host genes and viral genes/specific DVGs were calculated relative to GAPDH by RT-qPCR. *p<0.05, **p<0.01 by two-way ANOVA with Holm-Sidak’s multiple comparisons test. N=4, mean±SD (**C**) Supernatants from mock, DVG-B, SARS-CoV-2 and DVG-B/SARS-CoV-2 co-infected cells were collected and secreted IFNB1 was measured by ELISA. N=3, mean±SD. (**D**) Progeny infectious virus production was determined from supernatants of SARS-CoV-2, DVG-B/SARS-CoV-2 and DVG-A/SARS-CoV-2 co-infected cells **(A-B)** by TCID_50_/mL. Viral titration was performed in VeroAT cells. *p<0.05 by paired one-tailed t-test. N=4, mean±SD. (**E-F**) Cells were infected with SARS-CoV-2 (MOI 0.1) with increasing ratios of DVG-B: SARS-CoV-2 (**E**, MOI 0.1, 0.5, 2.5: MOI 0.1) or DVG-A: SARS-CoV-2 (**F**, MOI 0.1, 0.2, 0.5, 1: MOI 0.1). IFNB1, N and specific DVG copy numbers were quantified relative to GAPDH by RT-qPCR. SARS-CoV-2 titers, with increasing input MOIs of DVG-B or DVG-A, were measured by TCID_50_/mL. *p<0.05, **p<0.01, ***p<0.001 by one-way ANOVA with Dunnett’s multiple comparisons test. N=2, mean±SD.

To further compare the impact of DVG and WT virus infections on the cellular transcriptome, bulk RNA-seq was performed on infected cells in Fig. 4A-B. Principal component analysis (PCA) analysis separated DVG-B infection from mock, DVG-A, and WT virus infections, while DVG-A–infected cells exhibited a gene expression profile more similar to mock controls (Fig. S2A). First, comparison of DVG-B and WT virus infections identified over 10,000 differentially expressed genes (DEGs) (heatmap of all DEGs shown in Fig.S2B). These genes were grouped into 4 clusters. The majority of DEGs were upregulated in WT virus, but not DVG-B (cluster 1). Gene ontology (GO) analysis revealed that these genes were primarily enriched in TNFα-NFκB signaling and G protein coupled receptor signaling (Fig. 5A). 133 genes were downregulated in WT and WT+DVG-B and these genes were associated to endoplasmic reticulum stress and metabolic pathways (Fig. 5B). Surprisingly, only 69 genes were upregulated, while 29 genes were downregulated, in DVG-B–infected cells compared to mock infected controls. Consistent with our RT-qPCR results, antiviral response pathways were the most significantly enriched among genes upregulated in DVG-B infected cells (Fig. 5D), whereas no pathways were significantly enriched among the genes downregulated in DVG-B infected cells (Fig. 5C). We next performed similar comparisons to DVG-A vs. WT virus. Likewise, the majority of DEGs were associated with WT virus infection, and gene expression in co-infected cells was markedly reduced compared with WT virus infection alone, indicating that DVG-A significantly suppressed WT viral infection (Fig. S2C). Consistent with the PCA results, very few genes were differentially expressed specifically in DVG-A–infected cells (Fig. S2D). Taken together, our data directly demonstrated that both DVG-A and DVG-B suppressed WT virus replication. While the WT virus profoundly altered host transcriptome, DVG-B, but not DVG-A, selectively upregulated type I and type III antiviral responses.

**Fig 5.**
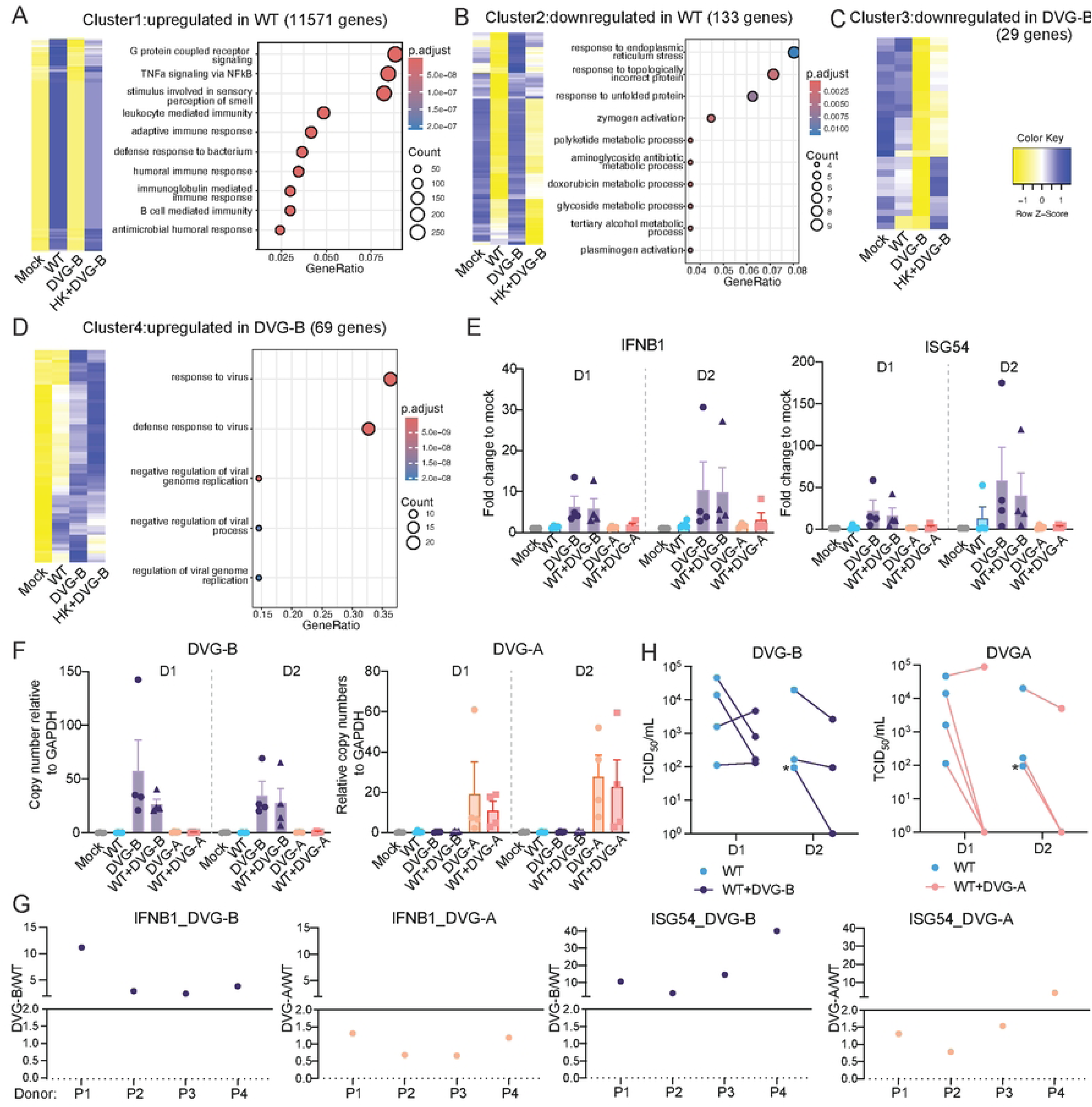
Transcriptomic profiling of WT SARS-CoV-2– and DVG-infected cells, and the effects of DVG-A and DVG-B infection in PCLS. (**A-D**) Total RNA from infected A549 cells was subjected to bulk RNA sequencing. Differentially expressed genes (DEGs, ≥twofold and ≤ 1% false discovery rate) were identified by comparing DVG-B–infected cells with mock-infected and WT SARS-CoV-2–infected cells. DEGs were grouped into four clusters, with heatmaps shown on the left and enriched pathways shown on the right: (A) cluster 1, (B) cluster 2, (C) cluster 3, and (D) cluster 4. Detailed DEG lists for each cluster were shown in Table S5. (**E-G**) PCLS were either singly infected or co-infected with WT virus and DVG-A or DVG-B. Co-infections were performed using a WT:DVG-B ratio of 1:5 and a WT:DVG-A ratio of 1:20. Supernatants and tissue slices were harvested at 24 and 48 h post infection. The expression of IFNB1 (**E**), ISG54 (**E**), DVG-B (**F**), and DVG-A (**F**) were examined in PCLS via qPCR. N=4, mean±SD. (**G**) Ratio of gene expression in DVG-B or DVG-A and WT virus infected tissue from all 4 donors (P1-P4). Higher ratio (>1) indicates that DVGs induced a higher gene expression than WT virus. (**H**) showing the infectious virus titer from supernatants via TCID_50_. Each dot represented one individual donor. * indicated that there were two donors with the same viral titers.

### DVG-B particles stimulated IFN responses in *ex vivo* human PCLS

To further assess whether DVG-B induces IFN responses in a physiologically relevant system, human PCLS from four adult donors were infected with DVGs and WT virus. Previous studies have shown that WT virus titer in PCLS peaks between 1 and 2dpi[34]; therefore, these time points were selected for analysis of IFN responses and infectious virus titers. We observed that DVG-B infection consistently elicited higher type I IFN responses compared with mock controls at both 1 and 2dpi, whereas this trend was not observed following DVG-A infection (Fig. 5E and 5F). Co-infection with WT virus and either DVG-A or DVG-B resulted in reduced viral titers in all four donors at 2dpi (Fig. 5H), further supporting the suppressive effects of both DVGs on WT virus replication. Interestingly, despite donor variability in responsiveness, the ratios of DVG-B to WT virus for IFNB1 expression (4/4 donors) and ISG54 expression (3/4 donors) were higher than those observed for DVG-A relative to WT virus (Fig. 5G), further indicating that DVG-B, but not DVG-A, robustly induced type I IFN responses *ex vivo*.

### DVG-B-derived dsRNAs displayed a distinct cellular distribution and were associated with increased IRF3 nuclear translocation than WT virus

Double stranded RNA (dsRNA) is the genomic replication intermediate during coronavirus infection and can be recognized by MDA5 or RIG-I to trigger innate IFN responses[21, 22]. However, dsRNA is normally sequestered within double membrane vesicles (DMV), also known as replication organelles (RO), which limits their detection via RLR pathways. We first examined the amount and cellular distribution of DVG-B-derived dsRNA by IFA and found that total dsRNA levels were significantly lower during DVG-B infection compared to WT (Fig. 6A; quantification in 6B), indicating reduced DVG-B replication levels. This is consistent with the markedly lower spike expression observed in DVG-B infected cells compared with WT virus (Figs. 3F and 6C). Interestingly, we found that while dsRNAs in WT infected cells were concentrated at perinuclear region, DVG-B-derived dsRNA was more diffusely distributed throughout the cell. This distinct distribution became apparent at 12 hpi and was prominent by 24 hpi (Fig. 6A), coinciding with the time point at which DVG-B infection significantly induced IFNB1 expression (Fig. 6C). We next co-stained DVG-B and WT infections for IRF3 with dsRNA at 24h post-infection. Consistently, despite fewer dsRNA positive cells, DVG-B infection resulted in a higher proportion of dsRNA+ cells exhibiting nuclear IRF3 than WT virus infection (>50% vs <10%), suggesting DVG-B-derived dsRNA/RNAs activated IRF3 more efficiently than WT (white arrows in Fig. 6D left panel, quantification in Fig. 6E). Because dsRNA formation requires viral genomic replication, dsRNA should not be detected when viral replication is inhibited. Therefore, we treated A549-ACE2 cells with remdesivir, an inhibitor of the SARS-CoV-2 RNA-dependent RNA-polymerase, before and during infection with DVG-B or WT virus. In the presence of remdesivir, no dsRNA was detected and IRF-3 remained majorly cytoplasmic at 24h post-infection (Fig. 6D, right panel). Correspondingly, DVG-B induced IFN production was completely abolished (Fig. 6F). To verify that remdesivir does not suppress virus entry, we infected remdesivir treated cells with DVG-A particles. Because DVG-A cannot replicate its genome independently, the amount of DVG-A detected from infected cells is the number of DVG-A particles that have entered the cells. As expected, remdesivir did not reduce DVG-A genomic levels (Fig. 6G). To further confirm that DVG-B induced the RLR pathway, we infected MAVS knockout (KO) and Protein Kinase R (PKR) KO A549 cells with DVG-B. As expected, MAVS KO, but not PKR KO cells, completely diminished the IFN induction (Fig. 6H). Taken together, our data indicated that the replication products of DVG-B, likely dsRNA, activated the RLR pathway to stimulate strong IFN responses.

**Fig 6.**
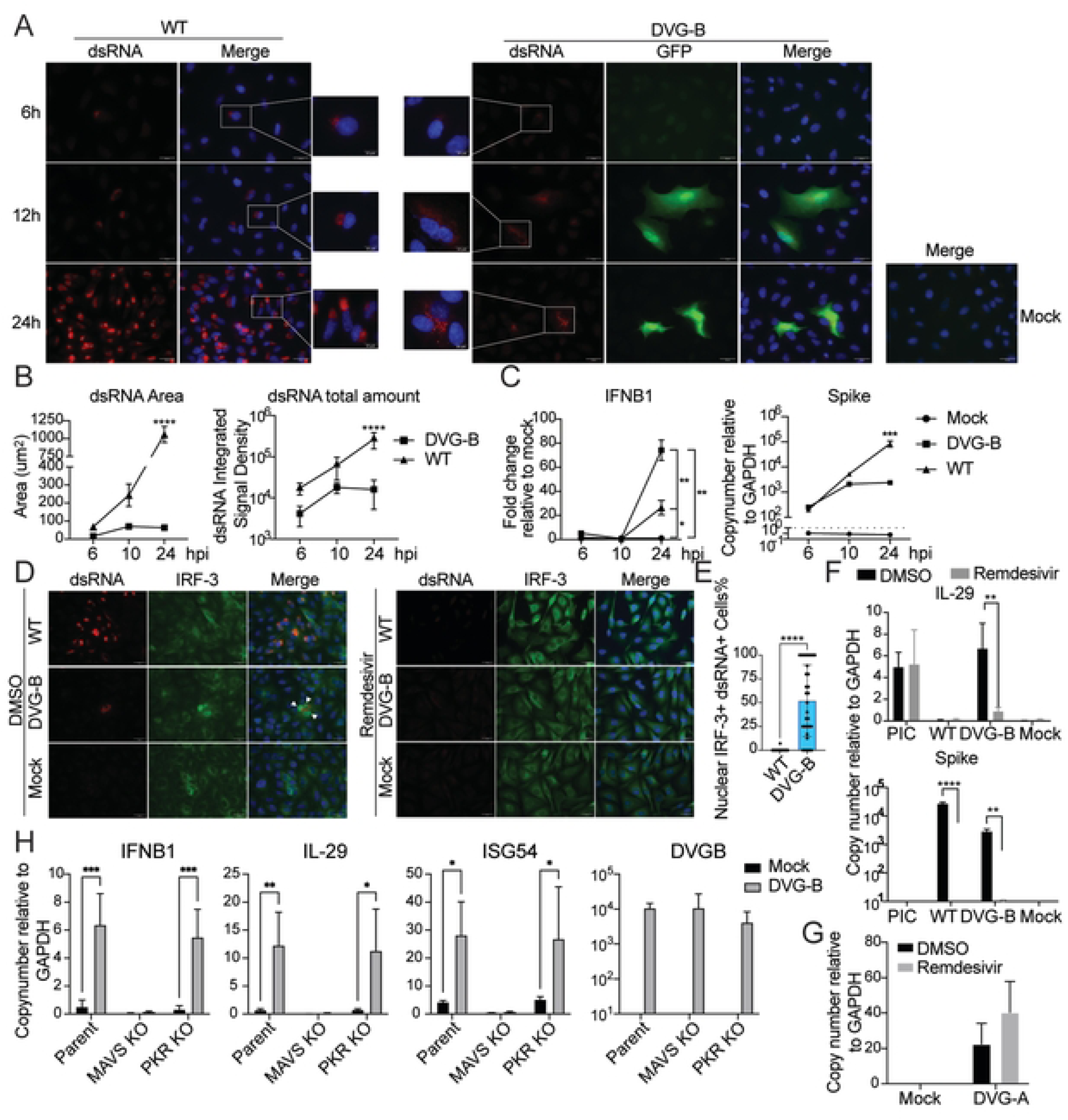
DVG-B derived dsRNA localizes to discrete puncta and induces significantly greater IRF-3 nuclear translocation compared to SARS-CoV-2. **(A-C)** A549-ACE2 cells were infected with SARS-CoV-2 or DVG-B (both MOI 1) and cells were collected at 6-, 10- and 24-hours post-inoculation for epifluorescence microscopy to (**A**) compare the cellular distribution of SARS-CoV-2 and DVG-B derived dsRNA or (**B**) quantify the total area of dsRNA signal and total dsRNA signal (integrated signal density). Insets depict magnified images of cells indicated by the white boxes. Scale bars indicate 10μm and 25μm, respectively. ****p<0.0001 by two-way ANOVA with Bonferroni’s multiple comparisons test. 6hpi N=4 fields. 10hpi N=5 fields. 24hpi, SARS-CoV-2 N=16 fields, DVG-B N=34 fields. Mean±SD. (**C**) IFNB1 and Spike gene expression were calculated relative to GAPDH by RT-qPCR. IFNB1 copy number values were normalized to mock for each respective timepoint. *p<0.05, **p<0.01, ***p<0.001 by two-way ANOVA with Turkey’s multiple comparisons test. N=3, mean±SD. (**D-E**) A549-ACE2 cells treated with 10μM remdesivir or an equal volume of DMSO were infected with SARS-CoV-2 or DVG-B (MOI 1) for 24-hours post-inoculation, respectively. N=2 biological repeats (**D**) Epifluorescence images of dsRNA and IRF-3 staining in DMSO (left) and remdesivir (right) treated cells. Scale bars indicate 25μM. Arrowheads indicate dsRNA+ cells with nuclear IRF-3. Remdesivir treated SARS-CoV-2 and DVG-B infections N=3 fields, respectively. DMSO treated SARS-CoV-2 infections N=14 fields, DVG-B infections N=30 fields. (**E**) Quantification of the proportion of dsRNA+ cells with nuclear IRF-3 relative to the total number of dsRNA+ cells per field. Each point represents one field. ****p<0.0001 by unpaired two-tailed t-test. SARS-CoV-2: 412 dsRNA+ cells among N=14 fields. DVG-B: 120 dsRNA+ cells among N=30 fields. (**F**) A549-ACE2 cells treated with 10μM remdesivir or an equal volume of DMSO were infected with SARS-CoV-2 or DVG-B (both MOI 1) or were transfected with low molecular weight poly(I:C) (PIC) for 24-hours post-inoculation, respectively. IL-29 and Spike gene copy numbers were calculated relative to GAPDH by RT-qPCR. **p<0.01, ****p<0.0001 by mixed-effects analysis with Bonferroni’s multiple comparisons test. N=3, mean±SD. (**G**) A549-ACE2 cells were treated with 10μM remdesivir or an equal volume of DMSO and infected with DVG-A (MOI 1) for 24-hours post-inoculation. DVG-A levels were quantified relative to GAPDH by RT-qPCR. N=4, mean±SD. (**H**) A549-ACE2 MAVS KO, PKR KO, or empty vector transduced control cells (parent) were infected with DVG-B (MOI 5) for 24-hours post-inoculation. IFNB1, IL-29, ISG54 and DVG-B copy numbers were quantified relative to GAPDH by RT-qPCR. *p<0.05, **p<0.01, ***p<0.001 by two-way ANOVA with Bonferroni’s multiple comparisons test. N=4, mean±SD.

To examine whether there were additional mutations and deletions in DVG-B stocks, we performed RNA deep sequencing on P2 and P3 DVG-B stocks, aligned RNA-seq reads to either the WT virus or DVG-B reference genomes, and visualized aligned files with IGV (Integrative Genomics Viewer) [35]. The introduced deletion at 27,700-29,674 was observed when aligned to WT viral reference genome, whereas this deletion was not observed in the alignment using DVG-B as the reference genome (Fig. S3A-B). Additional mutations were identified compared to the published sequences of CPER constructs (Table S2). Among them, 9 substitution mutations (8 nonsynonymous) were considered major, given that these mutations comprised more than 50% of the ribonucleotides at those positions (* in Table S2). Among the eight nonsynonymous substitutions, one occurred in nsp1 (no known function), one occurred in spike (adaptive mutation at S1/S2 cleavage site to enhance entry in Vero cells[36]), and the remaining seven were located in nsp12 (no known function). Based on the nsp12 crystal structure, the seven substitutions were clustered adjacent to nsp12 catalytic motif A[37]. Therefore, we introduced these mutations in nsp12 and further examined their effects on viral polymerase activity via a mini-replicon system[38]. This system is comprised of three helper plasmids expressing all 16 nonstructural proteins that form the SARS-CoV-2 viral polymerase complex (Vp), along with a backbone (BKB) plasmid containing the 5′UTR, a firefly luciferase reporter driven by the M gene TRS-B sequence, and the 3′UTR. A previous study demonstrated that firefly luciferase expression occurs only in the presence of the helper plasmids[38]. HEK293T cells were co-transfected with the BKB plasmid, a Renilla luciferase plasmid (transfection control), and either the WT Vp or nsp12 mutant Vp expressing plasmid. No significant difference in the firefly-to-Renilla luciferase activity ratio was observed between the WT and nsp12 mutant Vps (Fig. S3F), indicating that these nsp12 mutations do not substantially affect viral polymerase activity and are therefore unlikely to account for the altered dsRNA distribution.

In addition to point mutations, we identified additional deletions in the DVG-B stocks using the same pipeline[17]. These deletions were present at low levels in the P2 stock and became more enriched by P3 (Fig. S3C). The junctions were primarily located within ORF6 and the inserted GFP gene, near the 3′ end of the DVG-B genome (Fig. S3D). To determine whether these deletions contributed to DVG-B’s IFN induction phenotype, we infected A549-ACE2 cells with P2 and P3 DVG-B stocks at MOI 1. Both P2 and P3 DVG-B induced comparable IFN responses (Fig. S3E), indicating that these additional deletions did not alter DVG-B’s ability to induce IFNs. Therefore, the deep sequencing analyses suggests that the absence of the viral proteins encoded between positions 27,700 and 29,674 (the deletion range of hotspot B) were likely the major determinant of DVG-B’s distinct dsRNA distribution and robust IFN induction.

### Trans-complementation of the SARS-CoV-2 N protein partially restores the DVG-B-derived dsRNA pattern to the WT dsRNA phenotype, but not IFN phenotype

The SARS-CoV-2 nucleocapsid (N) protein is required for ribonucleocapsid formation and viral particle assembly. Additionally, multiple other key functions have been described for the N protein including undergoing liquid phase separation, interacting with the DMV via nsp3a, and innate immune antagonism[39–42] (cite). Therefore, to determine whether N complementation alters DVG-B dsRNA distribution, VeroAT or VeroE6:N cells were infected with DVG-B for 24 h. Confocal microscopy showed that DVG-B–derived dsRNA was widely distributed in VeroAT cells, similar to our observations in A549-ACE2 cells (Fig. 6A and S4A). In contrast, in VeroE6:N cells, most dsRNA localized to the perinuclear region and was closely associated with N (Fig. S4A). It should be noted that although N protein complementation dramatically altered the distribution of DVG-B-derived dsRNA, regions of dsRNA signal with no apparent colocalizing N protein were more abundant than in WT virus infected VeroE6:N cells.

Because Vero cells are deficient in type I IFN responses[43], we generated two A549-ACE2 cell lines constitutively expressing the SARS-CoV-2 N protein. The first expressed a strep-II tagged N-protein (A549:N) while the second lacked a protein tag (A549:N:NPT). A cell line transduced with an empty lentiviral vector encoding the puromycin resistance gene (A549:EV) was generated as a control. In A549:EV cells, WT-derived dsRNA localized to the perinuclear region and colocalized with N (R = 0.67; Fig. 7C, upper panel), whereas DVG-B–derived dsRNA remained dispersed (Fig. 7A). N complementation increased dsRNA levels and largely restored a WT-like distribution with N–dsRNA colocalization (Fig. 7A, quantification in 7C upper panel). Similar results were observed in A549:N:NPT cells, indicating that the strep-II tag did not affect dsRNA redistribution (Fig. S4D). Next, we examined the location of dsRNA in relation to DMVs. We used Nsp6 as an approximation for DMV localization given that Nsp6 zippered endoplasmic reticulum membranes reportedly connect to DMVs[44]. In WT infection, dsRNA strongly colocalized with Nsp6 in the perinuclear region (R = 0.74; Fig. 7B, quantification in Fig. 7C lower panel). During DVG-B infection in A549:EV cells, the Nsp6 signal was reduced and its correlation with dsRNA was reduced (R = 0.41). N complementation increased Nsp6 abundance and partially restored the dsRNA–Nsp6 association (R = 0.53, Fig. 7B), though not to WT levels. N also strongly correlated with Nsp6 during DVG-B infection in A549: N cells (R = 0.76, Fig. S4C). Taken together, our confocal data indicate that N complementation largely restored DVG-B derived dsRNA to the WT virus distribution pattern and suggest that the N protein influences DMV organization or maintenance. It should be noted that in both A549:N and VeroE6:N cells, dsRNA and Nsp6 patterns did not fully recapitulate the WT phenotype.

**Fig 7.**
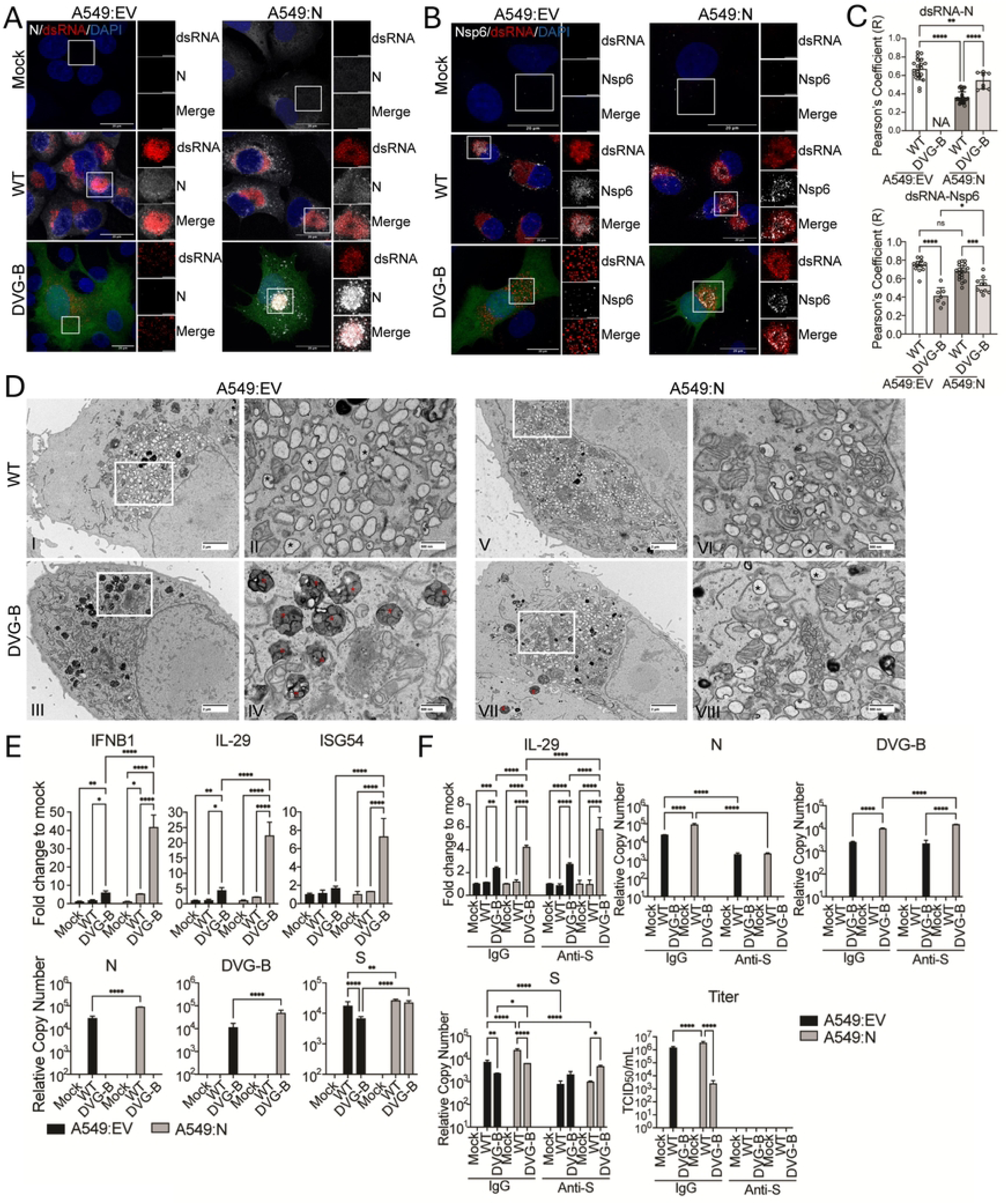
Trans-complementation of the SARS-CoV-2 N protein partially restores the DVG-B derived dsRNA pattern to the SARS-CoV-2 dsRNA phenotype. (A-B) A549:EV and A549:N cells were infected with WT (MOI 1) or DVG-B (MOI 1), respectively, and cells were collected 24-hours post-inoculation for confocal microscopy. **(A-B)** Maximum intensity projections of merged dsRNA (**A-B**, red) staining with the N protein (**A**, white) or Nsp6 (**B**, white) in A549-EV and A549:N cells. Nuclei were Hoechst stained (blue). Note that DVG-B encoded GFP is depicted (green). Scale bar indicates 20μm. Insets depict magnified cell regions indicated by the white boxes of dsRNA and N protein co-staining (**A**) or dsRNA and Nsp6 co-staining (**B**). Scale bar indicates 5μm. Graphs indicate the Pearson’s correlation coefficient (*R)* values for the co-localization of dsRNA and N (**C,** above) or dsRNA and Nsp6 (**C,** below) in A549:EV and A549:N cells, respectively. Individual values are shown. *p<0.05, **p<0.01, ***p<0.001, ****p<0.0001 by one-way ANOVA with Turkey’s multiple comparisons test. Mean±SD. (**D**) A549:EV (left) and A549:N cells (right) were infected with WT (MOI 1) or DVG-B (MOI 50 to maximize the DVG-B infected cell number), respectively, and cells were collected 24-hours post-inoculation for fixation and processing for transmission electron microscopy. Scale bars indicate 2μm. Insets depict magnified cell regions indicated by the white boxes. Scale bars indicate 500nm. Black asterisks indicate single membrane vesicles, red asterisks indicate lysosome-like structures. (**E**) Due to significant more cell death in WT infected A549:EV and A549:N cells, we reduced MOI of WT virus to 0.1 for this experiment. A549:EV and A549:N cells were infected with WT (MOI 0.1) or DVG-B (MOI 1), respectively, and cells were collected 24-hours post-inoculation for RT-qPCR. Expression of host genes and viral genes/DVG-B were calculated relative to GAPDH. IFNB1, IL-29 and ISG54 relative copy number values were normalized to mock for A549:EV and A549:N, respectively. Because the A549:N cell line expresses codon-optimized N, the qPCR primers specifically detect virus-derived N transcripts. *p<0.05, **p<0.01, ***p<0.001, ****p<0.0001 by two-way ANOVA with Turkey’s multiple comparisons test. N=2 biological repeats, mean±SD. (**F**) A549:EV and A549:N cells were inoculated as in (**E**). Immediately following 2-hours of inoculation, 10μg/mL of a neutralizing SARS-CoV-2 spike antibody or 10μg/mL of an IgG isotype control antibody were added to the infected cell cultures. 24-hours post inoculation cell supernatants were collected for virus titer determination by TCID_50_/mL quantification. Cells were collected for RT-qPCR. Expression of host genes and viral genes/DVG-B were calculated relative to GAPDH. IL-29 relative copy number values were normalized to mock for A549:EV and A549:N, respectively. *p<0.05, **p<0.01, ***p<0.001, ****p<0.0001 by two-way ANOVA with Turkey’s multiple comparisons test. N=3 biological repeats, mean±SD.

To further assess intracellular membrane rearrangements during DVG-B infection, transmission electron microscopy (TEM) was performed on WT virus– and DVG-B–infected A549:EV and A549:N cells, respectively (Fig. 7D). At 24 h post WT infection, numerous single-membrane vesicles (SMVs; indicated by *) were observed in infected cells of both cell types. Some of those SMVs contained virus particle-like structures, suggesting that they are likely the sites of viral particle assembly. In contrast, DVG-B–infected A549:EV cells showed a marked reduction in SMV abundance, suggesting that DVG-B is defective in the formation of the virus particle assembly site. Instead, these cells contained multiple large electron-dense membrane-bound structures with internal substructures, consistent with descriptions of tertiary lysosomes (indicated by * in Fig. 7D). In DVG-B–infected A549:N cells, however, SMV formation was partially restored. Notably, although tertiary lysosome–like structures were predominantly observed in DVG-B–infected A549:EV cells, they were also detected, albeit less frequently, in DVG-B–infected A549:N cells. Interestingly, we did not observe SMVs or lysosome-like vacuoles in mock-infected cells of either cell type, strongly suggesting that N is required, but independently insufficient, to induce SMV formation. Together, our TEM data indicate that DVG-B is defective in SMV formation, which can be partially rescued by N complementation.

To determine whether N complementation alters DVG-B–induced IFN responses, A549:EV and A549:N cells were infected with WT virus or DVG-B. RT-qPCR analysis revealed that in A549:EV cells, DVG-B induced higher IFNB1 and IL-29 expression than WT virus in both cell types, whereas WT virus triggered minimal antiviral responses, similarly as before (Fig. 7E). Unexpectedly, IFN responses were further increased in A549:N cells during DVG-B infection compared to A549:EV cells, correlating with increased viral RNA levels. Because complementation of the N protein to DVG-B infection now allows for virus particle formation, we hypothesized that the increase in IFN induction by DVG-B may be due to the formation and dissemination of infectious DVG-B particles. To address this, following 2-hours of SARS-CoV-2 or DVG-B inoculation, a spike neutralizing antibody (Anti-S) or IgG isotype control was added to infected cell cultures to prevent virus particle dependent spread. Similarly, we observed elevated IFN responses in A549:N cells compared to A549:EV cells at 24 h post DVG-B infection in the presence of IgG (Fig. 7F). As expected, neutralization successfully blocked the infectivity of virus particles in supernatants from both WT and DVG-B in A549:N cells (Fig. 7F, Titer). However, it did not reduce IFN induction or viral RNA levels in DVG-B–infected cells (Fig. 7F), suggesting that virus particle dissemination minimally contributes to the overall DVG-B RNA level even in the presence of N at 24hpi. In contrast, for WT virus, neutralizing antibody treatment significantly reduced the levels of N and S in both cell lines, while IL-29 levels remained similar to mock. Overall, these data indicate that, in our system, complementation of the N protein alone was sufficient to remodel the distribution of dsRNA/DMVs and restore the formation of SMVs during DVG-B infection. Despite these changes, N complementation was insufficient to reduce DVG-B induced antiviral gene expression to a level similar to that of WT virus.

## Discussion

As DVG populations are heterogeneous and have multiple roles in viral pathogenesis, it is suspected that different DVG species may exert distinct functions during infection. In this study, we demonstrate that DVGs generated from different regions of the SARS-CoV-2 genome exhibit distinct capacities to stimulate IFN responses. Despite the fact that both DVGs suppressed WT virus replication when supplemented at the onset of infection; DVG-A, generated from hotspot A, did not induce IFN responses, whereas DVG-B, generated from hotspot B, triggered robust IFN induction. Our data suggests that the IFN stimulation by DVG-B requires its genomic replication. In contrast, DVG-A’s inability to induce the IFN response is likely because it lacks the majority of the viral genome and therefore cannot independently replicate its genome. Due to this replicative defect, the abundance of DVG-A following supplementation was consistently lower than that of DVG-B, raising the possibility that insufficient DVG-A levels may account for the lack of IFN induction. To address this, we infected A549-ACE2 cells with DVG-A at an MOI of 80, exceeding the level of DVG-B detected in PCLS experiments, yet still did not observe any IFN stimulation compared to mock (Fig. S4E). Consistently, during *in vitro* viral infection (Fig. 2), cells positive for DVGs in hotspot A had a comparable level of DVGs compared to cells positive for DVGs in hotspot B, yet they exhibited a much lower level of IFN expression. Taken together, these findings strongly suggest that DVGs generated from hotspots A and B differ in their ability to stimulate IFN responses and therefore may play distinct roles in SARS-CoV-2 pathogenesis during natural infection. Further *in vivo* studies are required to validate this hypothesis.

Our RNA-seq analysis of virus- and DVG-infected cells revealed that DVG-B infection alone specifically induces antiviral responses, whereas WT virus infection broadly alters host transcriptional programs, prominently activating TNFα–NF-κB signaling pathways that may contribute to excessive inflammation. We speculate that this difference is due to the limited replication capacity of DVG-B, which appears sufficient to stimulate RLR pathways for IFN induction, but insufficient to trigger broader inflammatory pathways. For example, DVG-B infection does not produce infectious viral particles and generates only limited amounts of viral proteins and cellular membrane rearrangements compared with WT virus infection. In contrast, DVG-A infection minimally changed the host transcriptional responses compared with mock infected cells, further supporting the idea that DVG-A’s inhibition of WT virus occurs through an IFN-independent mechanism. This observation is consistent with a previous study in which a synthetic DVG with a genomic structure similar to DVG-A was developed as a therapeutic interfering particle (TIP) and shown to effectively suppress WT SARS-CoV-2 infection in Syrian golden hamsters without inducing IFN responses[45].

We observed that DVG-B-derived dsRNA was distributed throughout the cytoplasm, whereas WT virus derived dsRNA was primarily clustered in the perinuclear region. We initially hypothesized that this difference in dsRNA localization contributed to the robust IFN response induced by DVG-B. Complementation of the N protein during DVG-B infection partially restored dsRNA distribution to a pattern resembling that observed in WT virus infection and redistributed Nsp6 and N into close proximity with dsRNA. TEM further showed partial restoration of SMVs in N-expressing cells following DVG-B infection. However, this restoration did not reduce IFN expression to levels observed during WT virus infection. Instead, N complementation further enhanced IFN responses induced by DVG-B compared with infection in the absence of N. Together, these data indicate that while N plays an important role regulating dsRNA distribution, its complementation is not the major driver for limiting IFN induction during DVG-B infection. DVG-B’s other missing viral proteins may play important roles. Orf7a/7b and orf9b are all reported to target key factors downstream of RLR sensing in transfection-based systems. For example, orf7a was shown to reduce the expression level of TBK1, whereas orf7b was shown to interfere with the MAVS-TRAF6 interaction, which is essential to activate downstream IKK and TBK1[46]. Orf9b was also shown to block TBK1/IKK recruitment by blocking the Tom70/Hsp90 interaction[47]. Future studies with viruses specifically deleting orf7a, orf7b, orf9b, or N will assist in clarifying the individual contributions of these viral IFN antagonists to the robust IFN stimulation observed during DVG-B infection. Finally, the observation that N supplementation further enhances IFN responses suggests that, in addition to dsRNA, other viral RNA—likely viral genomic RNA—may serve as additional triggers of the strong IFN responses during DVG-B infection. Potentially, these viral RNAs are further amplified in the presence of N, thereby leading to increased IFN expression.

The large lysosome-like vacuoles observed in DVG-B–infected cells are intriguing. We speculate that these structures may arise from DVG-B’s virus particle assembly defect. Previous studies have reported that WT SARS-CoV-2 utilizes lysosomal membranes to form viral assembly sites[48]. It is therefore possible that DVG-B attempts to establish assembly sites, but in the absence of N protein, these structures are unstable or dysregulated and instead develop into enlarged lysosome-like vacuoles. Based on their size and spatial distribution relative to our dsRNA immuno-staining, these lysosome-like vacuoles may contain dsRNA. However, further experiments, such as confocal microscopy co-staining with lysosomal markers and dsRNA, as well as Correlative Light and Electron Microscopy (CLEM) using dsRNA and lysosomal markers, will be required to confirm their identity. If these lysosome-like vacuoles indeed contain dsRNA, this suggests that DVG-B’s dsRNA remains within membrane-bound compartments. In that case, it would be particularly interesting to determine how DVG-B’s dsRNA is sensed by cytosolic RLRs, or whether other viral RNA species serve as the primary triggers of RLR activation during DVG-B infection. Additionally, given the conservation of N amongst coronaviruses, this data raises the question as to whether those MERS-CoV or SARS-CoV DVGs lacking the nucleoprotein show similar patterns of dsRNA spatial dysregulation and lysosome-like structure formation[49].

While our VeroE6:N and A549:N cell systems yielded considerable shifts in the distribution of DVG-B–derived dsRNA, we consistently observed that N complementation failed to fully recapitulate the SARS-CoV-2 dsRNA pattern. This likely reflects fundamental differences between constitutive N expression and viral infection. Our cell lines may produce non-physiological levels of the N protein and inherently lack the regulated temporal control of N expression that occurs during SARS-CoV-2 infection. Moreover, N is normally expressed in concert with other viral proteins that collectively coordinate replication organelle formation and membrane remodeling. Viral infection of a cell line with pre-existing levels of N may alter the efficiency of these coordinated viral processes limiting the ability of N complementation to fully restore the WT phenotype during DVG-B infection. This may also potentially explain why dsRNA and N colocalization was significantly reduced during SARS-CoV-2 WT virus infection in A549:N cells compared to A549:EV cells. For the same reason, the titer of DVG-B particles in A549:N cells was more than three logs lower than that of WT virus, suggesting suboptimal assembly during DVG-B infection even in the presence of N. Consequently, DVG-B particle release is slower and less efficient, making early DVG-B dissemination less dependent on virus particle production.

Finally, we analyzed the DVG population in a COVID-19 cohort with reported disease severity scores. We observed a weak positive correlation between DVG abundance and disease severity. This modest association may reflect limitations of the dataset, including a relatively small sample size, a single time point of sample collection, and a cohort skewed toward severe cases. Nevertheless, approximately 40% of the DVG population mapped to hotspot B, indicating that this region is the dominant site of DVG generation compared with other genomic locations in patients. DVG+ patients also exhibited significantly higher viral loads than DVG-patients, suggesting that these DVGs likely arise as a consequence of high WT virus replication. Therefore, it is possible that hotspot B DVGs are generated relatively late during infection and their IFN-stimulating activity arises too late to effectively suppress WT virus replication. Instead, their presence may further amplify the inflammatory environment associated with high viral loads, thereby contributing to more severe disease outcomes. Larger, longitudinal cohorts will be necessary to better define the relationship between *de novo* DVG generation and COVID-19 severity.

In summary, our study provides direct evidence that SARS-CoV-2 DVGs generated from two distinct genomic regions exhibit different capacities to stimulate IFN responses. These findings provide a foundation for future studies to further elucidate the roles of DVGs in coronavirus pathogenesis and offer new insights into how DVGs may be harnessed as antiviral therapeutics.

## Materials and methods

### Cells, precision cut lung slices and viruses

All cell lines were cultured at 37°C 5% CO_2_ in tissue culture media (TCM) containing Dulbecco’s Modified Eagle’s Media (DMEM, Gibco) with 10% fetal bovine serum (FBS), 1mM sodium pyruvate (Gibco), 2mM L-glutamate (Gibco) and 50μg/ml gentamicin (Gibco). A549-ACE2 cells were obtained from Dr. Ruth Serra-Moreno’s lab (URMC) and were cultured in TCM supplemented with 10μg/mL blasticidin (Gibco). VeroE6 cells constitutively expressing ACE2 and TMPRSS2 (VeroAT) were cultured in TCM supplemented with 10μg/mL puromycin. A549-ACE2: MAVS KO, PKR KO, and empty vector control cells were obtained from Dr. Susan Weiss[50], University of Pennsylvania, and were maintained at 37°C 5% CO_2_ with RPMI 1640 (Gibco catalog no. 11875-093) containing 10% FBS, 100U/ml penicillin, and 100μg/mL streptomycin. All cells were treated with mycoplasma removal agent before experimentation (MP Biomedicals). Human precision cut lung slices (PCLS) were obtained from LungMAP (URMC) and were processed according to protocols published elsewhere[51, 52]. All four PCLS donors were between 20-60 years of age. PCLS were maintained at 37°C 5% CO_2_ in DMEM containing 10% FBS, 1 mM sodium pyruvate, 1X non-essential amino acids, 15 mM HEPES with 0.25μg/mL of amphotericin B (Gibco), and 80ng/mL gentamicin (Gibco). After 24 hours at 37°C, media was replaced and PCLS were incubated for an additional 24 hours prior to infection. SARS-CoV-2 hCoV-19/Hong Kong/VM20001061/2020 (HK, BEI NR-52282) was obtained through BEI resources. Viruses were propagated in VeroE6 cells in the presence of 5% FBS TCM. To limit the accumulation of DVGs, cells were infected at MOI 0.001 and virus containing supernatants were collected 3 days post-infection and filtered (0.45μm).

### DVG identification and association with IFN responses from bulk and scRNA-seq datasets

For Fig. 2B-2D, we used the publicly available dataset from Ravindra et al. 2021 accessed through the NCBI database (GSE166766). This study consisted of single cell RNA-Seq (scRNA-Seq) data from human bronchial epithelial cells (NHBEs) infected with SARS-CoV-2 that were harvested 1 day post infection (dpi), 2 dpi, and 3 dpi. The detailed approach for how we identify DVGs and DVG+ cells from scRNA-seq has been previously published elsewhere[17]. Briefly, we first used Cell Ranger [53] to construct gene expression matrices for each sample. To identify the number of viral transcripts, the SARS-CoV-2 reference sequence was concatenated to the end of the human genome reference as one additional gene. The gene expression matrices were then loaded into the Seurat package in R [54]. Cells were then clustered and annotated based on the gene makers used in the original publication of this dataset. To identify DVGs, we used UMI-tools [55] to associate the cell barcodes and UMIs with each corresponding read name. We used Bowtie2 [56], ViReMa, and a custom R filtering script for DVG identification. We then used the filtered ViReMa output to re-quantify DVG count based on the UMIs associated with each cell barcode. Specifically, we counted the number of unique UMIs with the same cell barcode as the total number of DVG reads per cell. The numbers of DVG UMIs of each cell barcode and the junction positions was then added to the gene expression matrix created by Cell Ranger. Cells with more than one DVG UMI (virus positive cells) were grouped as DVG+. Dot plots associating DVG junction positions and gene expression in DVG+ cells were graphed in R with the raw spread sheet containing junction positions and IFN expression shown in Table S4. Cells positive for DVG-A or DVG-B were defined based on the junction positions of the DVGs they contained. The boundaries of DVG-A and DVG-B were determined according to a previously published study[17]. For bioinformatic analysis of the human cohort (Fig. 1), we used the publicly available dataset (SCP1289[29]). This study consisted of scRNA-Seq data from a cohort study within known WHO severity scores (0-8) and age information. This cohort study consisted of nasal samples from 58 donors, 37 COVID-19 patients and 21 control patients. Within 37 COVID-19 patients, 23 individuals are less than 65 years of age (younger) and 14 elderly with age >65. We used similar approach illustrated above to identify DVGs from this scRNA-seq dataset. Patients with at least one unique DVG UMI were considered as DVG+ patients. The raw value of DVGs for individual patients were summarized in Table S3.

For Fig. 2E-2F, we used the publicly available dataset accessed through the NCBI database (GSE213759). This study consisted of bulk RNA-Seq data from Calu3 cells infected with 2000 E copies of different SARS-CoV-2 variants of concerns. Analyzed samples were infected cells collected at 24 – and 48-hpi. The detailed approach for how we identify DVGs from bulk RNA-seq has been previously published elsewhere[17]. Briefly, we first used Bowtie2 (v. 2.2.9, [56]) to align the reads to the GRCh38 human reference genome. The unmapped reads were then applied to ViReMa (Viral-Recombination-Mapper v. 0.21) to identify recombination junction sites and their corresponding read counts using SARS-CoV-2 reference genome (GenBank ID MT020881.1). A custom filtering script was developed in R to identify junction reads that met our criteria. We required the positions of both sites (break and rejoin) of junction reads larger than 85, as TRS-L is reported to be located with the first 85 nts of the SARS-CoV-2 genome. Additionally, we required deletions longer than 100 nts to ensure that the truncated viral RNAs are deficient in replication. We included all deletions with counts >10. The number of viral reads (reads fully aligned to WT viral reference genome) in each bulk RNA-Seq sample was quantified using the RSubread Bioconductor package. The junction frequency (J_freq_) was calculated by DVG counts over viral read counts.

### Generation of specific DVG particles and co-infection with WT virus

The DVG-A rejoin and break point boundaries are positions 700, located within the nsp1 gene sequence, and 29,674, within the 3’UTR, respectively. The 5′UTR–700 nt fragment (5′–700) and the 29,674–3′UTR fragment (29,674–3′) were amplified from CPER fragments[33] (kindly provided by Dr. Yoshiharu Matsuura of Osaka University). The mKate2 gene and the PS9 sequence (viral packaging signal; ref) were inserted between the 5′–700 and 29,674–3′ fragments by overlapping PCR. The final construct was cloned into the pcDNA3 vector. To generate SARS-CoV-2 virus like particles specifically packaging DVG-A, 3x10^6^ HEK293T cells were plated on poly-d-lysine (50μg/mL) treated 10cm dishes 1 day prior to transfection. Each dish was transfected with 3.35μg pCDNA3.1-M-IRES-E (addgene #177938), 6.65μg pCDNA3.1-N (addgene #177937), 0.25μg pCDNA3.1-S (addgene #177939), and 10μg pCDNA3.1-DVG-A at a 3:1(v/w) PEI (1mg/mL):DNA(μg) ratio. Transfection complexes were added dropwise and cells were incubated for 24 hours at 37°C. 1 day post-transfection, the transfection media was aspirated and 10% TCM without antibiotics was added and cells were incubated for 24 hours at 37°C. 2 days post-transfection cell supernatants were collected and quick frozen, and 10% FBS TCM without antibiotics was added and cells were incubated for 24 hours at 37°C. Supernatant collection proceeded for 5 days. DVG-A containing supernatants were filtered (0.45μm) and centrifuged at 5000 x g, 4°C, for 16 hours. The DVG-A containing pellet was resuspended in 5% TCM and ultracentrifuged using an SW-32 Ti rotor (Beckman Coulter) through a sterile filtered 30% sucrose cushion (50mM Tris-HCL, 100mM NaCl, 50mM maleic acid, 5mM EDTA, in nuclease free water, pH=6) at 100,000 x g, 4°C, for 2 hours. The DVG-A containing pellet was washed 5 times with HBSS prior to resuspension in 5% FBS TCM. The generation of DVG-B was adapted from CPER method to rescue recombinant SARS-CoV-2 infectious clones described by Torii et. al[33] and is derived from SARS-CoV-2/Hu/DP/Kng/19-020. Briefly, the ORF7a-N deletion found in DVG-B was introduced into gene fragment F9/10 by replacing the ORF7A-N genes with the enhanced green fluorescent protein (eGFP) gene under the N TRS-B sequence. To generate SARS-CoV-2 virus like particles specifically packaging DVG-B, per 6-well, half of the DVG-B CPER products (25 μl), together with different combinations of pCDNA3.1-M-IRES-E (0.67 μg), pCDNA3.1-N (1.32μg), or pCDNA3.1-S (0.02μg), were transfected into HEK293T cells using lipofectamine 2000 similarly as the approach used to package DVG-A. Supernatants were collected daily from days 2-8 post transfection. All supernatants were then pooled, filtered, and concentrated by centrifugation of 5000 x g 4°C, for 16 hours. Concentrated DVG-B was then serially passaged three times in VeroE6:N cells at MOI 0.001 for 5-6 days per passage. The working stock of DVG-B were purified via 30% sucrose cushion before usage.

For infection, cells were inoculated with DVG-A or DVG-B alone, or co-infected with SARS-CoV-2 plus DVG-A or DVG-B, for 3 h at 37 °C in 5% FBS tissue culture medium, using the MOIs and DVG-to-SARS-CoV-2 MOI ratios specified in the corresponding figure legends. Each PCLS was inoculated with 1x10^5^ TCID_50_ SARS-CoV-2, 5x10^5^ FFU DVG-B or 2x10^6^ genome copy numbers of DVG-A, either alone or in combination, in PCLS maintenance media (5% FBS) for 3 hr at 37°C. Post inoculation, PCLS were washed three times with DPBS and cultured in fresh PCLS maintenance medium (5% FBS). Both supernatants and tissue slices were harvested at 24h or 48h post-infection. To harvest PCLS, tissues were homogenized at 4°C in 1mL Trizol and total RNA was extracted.

### Virus and DVG Titration

SARS-CoV-2 titers were determined by median tissue culture infectious dose (TCID_50_). Briefly, VeroAT cells were infected with serially diluted viral supernatants and CPE was measured 3 days post-infection. TCID_50_/mL was quantified using the Reed-Muench method. DVG-B titers were calculated by determining the fluorescent forming units (FFU)/mL. Briefly, VeroE6:N cells were infected with serially diluted DVG supernatants and fluorescent cells were quantified 24 hours post-infection. To determine the DVG-A titer, total RNA was extracted from DVG-A stocks by Trizol-LS and purified RNA (2μg) was Turbo DNaseI (2U/μg RNA) treated at 37°C for 30 minutes. DVG-A copy number/mL was quantified relative to a standard curve of serially diluted DVG-A encoding plasmid by RT-qPCR.

### Remdesivir treatment

1 day prior to infection A549-ACE2 cells were treated with 10 μM remdesivir (MedChemExpress catalog no. GS-5734) or and equal volume of DMSO in 10% TCM. Cells were inoculated with SARS-CoV-2 HK (MOI 0.2), DVG-B (MOI 1) in 5% TCM for 3 hours in the absence of remdesivir or DMSO. Following inoculation, 5% TCM containing 10 μM remdesivir or an equal volume of DMSO was added to the cells. Alternatively, cells were transfected with 1 μg/mL low molecular weight poly(I:C) (InvivoGen catalog no. tlrl-picw) via lipofectamine 2000. 24 hours post-inoculation/transfection cells were lysed in Trizol for total RNA extraction or fixed in 4% paraformaldehyde for immunostaining. DVG-A (MOI 10) infections were performed identically followed by Trizol lysis for total RNA extraction.

### RNA extraction and RT-qPCR

Tissue culture cell lines/supernatants were lysed in TRIzol or TRIzol LS (Invitrogen) at the indicated timepoints and total RNAs were extracted according to the manufacturer’s specifications, respectively. Total RNA (100ng-1μg) was reversed transcribed using the high-capacity RNA to cDNA kit (Applied Biosystems). cDNA was diluted and qPCR reactions were performed in triplicate using specific RT-qPCR primers (Table S1) and Power Track SYBR Green PCR Master Mix (Applied Biosystems) in a Viia7 Light-cycler (Applied Biosystems). Relative copy numbers for viral and host gene expression levels were calculated by normalizing to the GAPDH copy.

### RNA-Seq of A549-ACE2 infections and DVG-B stocks and host transcriptome analysis

RNAs from virus and DVG infected A549-ACE2 cells were extracted, followed by DNase I digestion. 100-300 ng of total RNAs were used for single-end bulk RNA-seq (Plasmidsaurus). Quality of the fastq files was assessed using FastQC v0.12.1. Reads were then quality filtered using fastp v0.24.0 with poly-X tail trimming, 3’ quality-based tail trimming, a minimum Phred quality score of 15, and a minimum length requirement of 50 bp. For transcriptome analyses, quality-filtered reads were aligned to the human reference genome (hg38) using bowtie2. Gene-level expression quantification was performed using featureCounts (subread package v2.1.1) with strand-specific counting, multi-mapping read fractional assignment, exons and three prime UTR as the feature identifiers, and grouped by gene_id. Final gene counts were annotated with gene biotype and other metadata extracted from the reference GTF file. Sample-sample correlations for sample-sample heatmap and PCA were calculated on normalized counts (TMM, trimmed mean of M-values) using Pearson correlation. Differential expression was done with edgeR v4.0.16 using standard practice including filtering for low-expressed genes. Functional enrichment, when available, is performed using gene set enrichment analysis. Detailed differentiated gene lists for designated clusters were shown in Table S5-S9.

For P2 and P3 DVG-B stocks, total RNA was extracted using TRIzol reagent followed by Turbo DNaseI treatment. RNA quality was assessed using the RNA Pico 6000 module on an Agilent Tapestation 2100 (Agilent Technologies) prior to cDNA library preparation. 100 ng of total cDNA libraries were prepared using the Illumina TruSeq Stranded Total RNA LT kit with Ribo-Zero Gold, according to the manufacturer’s instructions. Samples were run on Illumina NextSeq 500 to generate 150 bp, paired-end reads, resulting in 20-50 million reads per sample, with an average Q30 score ≥ 96.8%. Reads were aligned to DVG-B reference genome and WT virus genome (MT020881). The resulting BAM files were used for visualization in IGV, and unmapped reads were analyzed using ViReMa for DVG identification, as described in the previous DVG identification section.

### Transduction of cells to constitutively express SARS-CoV-2 N

VeroE6 cells constitutively expressing the SARS-CoV-2 N protein (VeroE6:N) were generated by lentiviral transduction using the pLVX-EF1α-IRES-Puro (Takara) lentiviral expression vector followed by puromycin selection and single clone propagation. VeroE6:N cells were cultured in 10% FBS tissue culture media supplemented with 8 μg/ml puromycin. A549-ACE2 cells constitutively expressing SARS-CoV-2 N (A549:N) or an empty vector control (A549:EV) were similarly generated by lentiviral transduction followed by puromycin selection and single clone propagation. Lentiviral constructs were obtained from addgene (https://doi.org/10.1038/s41586-020-2286-9). A549:N and A549:EV cells were cultured in 10% TCM containing 10μg/ml blasticidin and 1.5μg/ml puromycin.

### Immunofluorescent assay, confocal microscopy, and image quantification

For widefield and confocal microscopy, at the indicated times post-infection, cells were fixed on #1.5 coverslips with 4% paraformaldehyde for 15 minutes at room temperature and washed 3 times with 1X PBS. Fixed cells were permeabilized and blocked in 1X PBS containing 0.3% Triton X-100 and 5% goat serum (Cell Signaling) for 1 hour at room temperature. Primary and secondary antibodies were diluted in 1X PBS containing 1% bovine serum albumin and 0.3% Triton X-100. Primary antibody incubations were carried out for 18 hours at 4°C. Secondary antibody incubations were carried out for 2 hours at room temperature. See Table S1 for antibody concentrations. Cells were then incubated with 1μg/mL Hoechst (Invitrogen H3570, 10mg/mL) for 10 minutes at room temperature. Cover slips were mounted onto microscope slides using the ProLong Gold Antifade mountant (Invitrogen). Widefield images were taken with a Leica DMIRB inverted fluorescence microscope with a cooled charge-coupled device (Cooke) on Image-Pro Plus software (Media Cybernetics). The proportion of dsRNA+ cells containing nuclear IRF-3 in SARS-CoV-2 and DVG-B infections was quantified by normalizing the number of dsRNA+nuclear IRF-3+ cells to the total number of dsRNA+ cells per image. Confocal microscopy images of WT and DVG-B infected A549:EV and A549:N expressing cells were captured using a Nikon A1R HD laser scanning confocal microscope with TIRF using a 60X/1.49 oil objective. 405nm, 488nm, 561nm and 640nm excitation diodes to excite Hoechst, eGFP, dsRNA/N and N/Nsp6, respectively. Confocal images of HK and DVG-B infected VeroAT and VeroE6:N expressing cells were captured using a Leica Stellaris 5 inverted confocal microscope with WLL using a 63X/1.40 oil objective. Widefield and confocal microscopy images were analyzed using Fiji/ImageJ. Pearson correlation (*R*) values were determined using the Coloc 2 Fiji plugin on the average intensity projections of z-stacks obtained by confocal microscopy.

### Transmission Electron Microscopy

Infected cells cultured on glass chamber slides were fixed in 0.1M Millonigs phosphate buffer containing 5% glutaraldehyde overnight at 4°C. The slides were washed 3X with 0.1M phosphate buffer, post-fixed for 20 minutes in buffered 1.0% osmium tetroxide/1.5% potassium ferrocyanide, dehydrated on the slides (chamber removed) through a graded series of ethanol (10-minute incubations) to 100%. Then 1:1 100% ethanol/Spurr’s resin for 15 minutes before being transferred to 100% Spurr’s epoxy resin twice, once for an hour and secondarily overnight. The next day size 3 BEEM capsules were filled with fresh Spurr’s resin and inverted and placed over the cells of interest on the slides. The slides/capsules were polymerized 24 hours at 65^0^C. The next day the slides were dipped several times in liquid nitrogen to break the surface tension between the glass and the polymerized capsules until they “popped off” the slide[57]. The capsules with the entrapped cells were trimmed down to a trapezoid and thin sectioned with a diamond knife at 70nm and placed onto 2x1 mm slotted formvar/carbon copper grids. The grids were stained with uranyl acetate and lead citrate and examined using a Hitachi (Pleasanton, CA) camera and an attached AMT Nanosprint (Woburn, MA) 12-megapixel digital camera.

### ELISA

IFNB1 concentrations within the supernatants of SARS-CoV-2, DVG-B and SARS-CoV-2/DVG-B co-infected cells were determined using the VeriKine-HS Human Interferon-Beta TCM ELISA Kit (PBL Assay Science catalog no. 41435) according to the manufacturer’s instructions. Absorbance at 450nm was quantified using a SpectraMax Mini multi-mode microplate reader. A 4-parameter logistic plot with 1/y^2^ weighted analysis was used for the results calculation.

### Neutralization assay

A549:EV and A549:N cells were infected with WT or DVG-B at designated MOIs similarly. Post inoculation, cells were washed three times with PBS and cultured in 5% TCM with 10μg/mL of Spike neutralizing antibody (40592-R001, Sino Biological, obtained from BEI resources). Both supernatants and infected cells were harvested 24 hr post infection.

### Statistical analysis

All statistical analyses were performed with GraphPad Prism version 5.0 (GraphPad Software, San Diego, CA) and *R* v*3.4.1*. A statistically significant difference was defined as a *p*-value <0.05. by one way or two way analysis of variance (ANOVA) with a *post hoc* test to correct for multiple comparisons, or Student’s t-test (based on specific data sets as indicated in each figure legend).

### Data availability

Bulk RNA-seq datasets for DVG-B and A549-ACE2 infected cells are in the process of depositing to the Gene Expression Omnibus (GEO) database for public access.

## Acknowledgements

We thank the Electron Microscopy Resource in the Center for Advanced Research Technologies at the University of Rochester Medical Center for processing the samples (RRID#: SCR_012366). We thank the URMC Center for Advanced Microscopy & Nanoscopy (RRID:SCR_023177). We thank Dr. Susan Weiss (University of Pennsylvania) for kindly providing the A549-MAVS-KO and A549-PKR-KO cell lines. We would also like to thank LungMAP for providing us the human precision-cut lung slices. Conceived experiments: J.W.B., and Y.S.; Developed methodology: J.W.B, and Y.S.; Performed experiments, collected and analyzed data: J.W.B, S.S., X.W., H.A., S.C., Y.S.; Provided reagents and resources: P.G., R.S.M., T.J.M., Y.S.; Wrote the original draft: J.W.B., and Y.S; Supervised research activities: R.S.M., T.J.M., Y.S.

## Supporting information captions

**S1 Fig. DVGs clustered in hotspot A and B showed different IFN induction ability via single-cell RNA-seq.**

**S2 Fig. Transcriptome profiling of WT virus and DVG infected A549-ACE2 cells.**

**S3 Fig. RNA sequencing analyses of DVG-B stocks.**

**S4 Fig. Validation of DVG-B’s dysregulated dsRNA distribution phenotype.**

**S1 Table. Oligo primer and antibody list.**

**S2 Table. Mutations identified in DVG-B stocks.**

**S3 Table. Total DVG abundance and individual species identified from Cohort study.**

**S4 Table. DVG junctions associated with their single-cell IFN expression at 2dpi.**

**S5-S9 Table. DEG lists of different Clusters from Fig.5**.

## References

1. Von Magnus P. Studies on Interferenee in Experimental Influenza. I. Biological Observations. Arkiv for Kemi, Mineralogi och Geologi. 1947;(7).

2. Li D, Lott WB, Lowry K, Jones A, Thu HM, Aaskov J. Defective interfering viral particles in acute dengue infections. PLoS One. 2011;6(4):e19447. Epub 20110429. doi: 10.1371/journal.pone.0019447. PubMed PMID: 21559384; PubMed Central PMCID: PMCPMC3084866.

3. Calain P, Monroe MC, Nichol ST. Ebola virus defective interfering particles and persistent infection. Virology. 1999;262(1):114–28. doi: 10.1006/viro.1999.9915. PubMed PMID: 10489346.

4. von Magnus P. Incomplete forms of influenza virus. Advances in virus research. 2: Elsevier; 1954. p. 59–79.

5. Calain P, Roux L. Generation of measles virus defective interfering particles and their presence in a preparation of attenuated live-virus vaccine. J Virol. 1988;62(8):2859–66. doi: 10.1128/jvi.62.8.2859-2866.1988. PubMed PMID: 3392771; PubMed Central PMCID: PMCPMC253722.

6. Treuhaft MW, Beem MO. Defective interfering particles of respiratory syncytial virus. Infect Immun. 1982;37(2):439–44. doi: 10.1128/iai.37.2.439-444.1982. PubMed PMID: 6288562; PubMed Central PMCID: PMCPMC347553.

7. Girgis S, Xu Z, Oikonomopoulos S, Fedorova AD, Tchesnokov EP, Gordon CJ, et al. Evolution of naturally arising SARS-CoV-2 defective interfering particles. Commun Biol. 2022;5(1):1140. Epub 20221027. doi: 10.1038/s42003-022-04058-5. PubMed PMID: 36302891; PubMed Central PMCID: PMCPMC9610340.

8. Tapia K, Kim WK, Sun Y, Mercado-López X, Dunay E, Wise M, et al. Defective viral genomes arising in vivo provide critical danger signals for the triggering of lung antiviral immunity. PLoS Pathog. 2013;9(10):e1003703. Epub 20131031. doi: 10.1371/journal.ppat.1003703. PubMed PMID: 24204261; PubMed Central PMCID: PMCPMC3814336.

9. Sun Y, Jain D, Koziol-White CJ, Genoyer E, Gilbert M, Tapia K, et al. Immunostimulatory Defective Viral Genomes from Respiratory Syncytial Virus Promote a Strong Innate Antiviral Response during Infection in Mice and Humans. PLoS Pathog. 2015;11(9):e1005122. Epub 2015/09/04. doi: 10.1371/journal.ppat.1005122. PubMed PMID: 26336095.

10. Johnston MD. The characteristics required for a Sendai virus preparation to induce high levels of interferon in human lymphoblastoid cells. J Gen Virol. 1981;56(Pt 1):175–84. doi: 10.1099/0022-1317-56-1-175. PubMed PMID: 6170730.

11. Marcus PI, Gaccione C. Interferon induction by viruses. XIX. Vesicular stomatitis virus--New Jersey: high multiplicity passages generate interferon-inducing, defective-interfering particles. Virology. 1989;171(2):630–3. doi: 10.1016/0042-6822(89)90637-5. PubMed PMID: 2474895.

12. Strähle L, Marq JB, Brini A, Hausmann S, Kolakofsky D, Garcin D. Activation of the beta interferon promoter by unnatural Sendai virus infection requires RIG-I and is inhibited by viral C proteins. J Virol. 2007;81(22):12227–37. Epub 20070905. doi: 10.1128/jvi.01300-07. PubMed PMID: 17804509; PubMed Central PMCID: PMCPMC2169027.

13. Yount JS, Gitlin L, Moran TM, López CB. MDA5 participates in the detection of paramyxovirus infection and is essential for the early activation of dendritic cells in response to Sendai Virus defective interfering particles. The Journal of Immunology. 2008;180(7):4910–8.

14. Shingai M, Ebihara T, Begum NA, Kato A, Honma T, Matsumoto K, et al. Differential type I IFN-inducing abilities of wild-type versus vaccine strains of measles virus. J Immunol. 2007;179(9):6123–33. doi: 10.4049/jimmunol.179.9.6123. PubMed PMID: 17947687.

15. Felt SA, Sun Y, Jozwik A, Paras A, Habibi MS, Nickle D, et al. Detection of respiratory syncytial virus defective genomes in nasal secretions is associated with distinct clinical outcomes. Nat Microbiol. 2021;6(5):672–81. Epub 2021/04/03. doi: 10.1038/s41564-021-00882-3. PubMed PMID: 33795879.

16. Xu J, Sun Y, Li Y, Ruthel G, Weiss SR, Raj A, et al. Replication defective viral genomes exploit a cellular pro-survival mechanism to establish paramyxovirus persistence. Nat Commun. 2017;8(1):799. Epub 20171006. doi: 10.1038/s41467-017-00909-6. PubMed PMID: 28986577; PubMed Central PMCID: PMCPMC5630589.

17. Zhou T, Gilliam NJ, Li S, Spandau S, Osborn RM, Connor S, et al. Generation and Functional Analysis of Defective Viral Genomes during SARS-CoV-2 Infection. mBio. 2023;14(3):e0025023. Epub 20230419. doi: 10.1128/mbio.00250-23. PubMed PMID: 37074178; PubMed Central PMCID: PMCPMC10294654.

18. Llanes A, Restrepo CM, Caballero Z, Rajeev S, Kennedy MA, Lleonart R. Betacoronavirus Genomes: How Genomic Information has been Used to Deal with Past Outbreaks and the COVID-19 Pandemic. Int J Mol Sci. 2020;21(12). Epub 20200626. doi: 10.3390/ijms21124546. PubMed PMID: 32604724; PubMed Central PMCID: PMCPMC7352669.

19. Perlman S, Netland J. Coronaviruses post-SARS: update on replication and pathogenesis. Nat Rev Microbiol. 2009;7(6):439–50. doi: 10.1038/nrmicro2147. PubMed PMID: 19430490; PubMed Central PMCID: PMCPMC2830095.

20. Magnen M, You R, Rao AA, Davis RT, Rodriguez L, Bernard O, et al. Immediate myeloid depot for SARS-CoV-2 in the human lung. Sci Adv. 2024;10(31):eadm8836. Epub 20240731. doi: 10.1126/sciadv.adm8836. PubMed PMID: 39083602; PubMed Central PMCID: PMCPMC11290487.

21. Yin X, Riva L, Pu Y, Martin-Sancho L, Kanamune J, Yamamoto Y, et al. MDA5 Governs the Innate Immune Response to SARS-CoV-2 in Lung Epithelial Cells. Cell Rep. 2021;34(2):108628. Epub 2021/01/14. doi: 10.1016/j.celrep.2020.108628. PubMed PMID: 33440148; PubMed Central PMCID: PMCPMC7832566.

22. Minkoff JM, tenOever B. Innate immune evasion strategies of SARS-CoV-2. Nat Rev Microbiol. 2023;21(3):178–94. Epub 20230111. doi: 10.1038/s41579-022-00839-1. PubMed PMID: 36631691; PubMed Central PMCID: PMCPMC9838430.

23. Hadjadj J, Yatim N, Barnabei L, Corneau A, Boussier J, Smith N, et al. Impaired type I interferon activity and inflammatory responses in severe COVID-19 patients. Science. 2020;369(6504):718–24. Epub 20200713. doi: 10.1126/science.abc6027. PubMed PMID: 32661059; PubMed Central PMCID: PMCPMC7402632.

24. Chaturvedi S, Vasen G, Pablo M, Chen X, Beutler N, Kumar A, et al. Identification of a therapeutic interfering particle-A single-dose SARS-CoV-2 antiviral intervention with a high barrier to resistance. Cell. 2021;184(25):6022–36.e18. Epub 2021/11/29. doi: 10.1016/j.cell.2021.11.004. PubMed PMID: 34838159; PubMed Central PMCID: PMCPMC8577993.

25. Rand U, Kupke SY, Shkarlet H, Hein MD, Hirsch T, Marichal-Gallardo P, et al. Antiviral Activity of Influenza A Virus Defective Interfering Particles against SARS-CoV-2 Replication In Vitro through Stimulation of Innate Immunity. Cells. 2021;10(7):1756. Epub 2021/08/08. doi: 10.3390/cells10071756. PubMed PMID: 34359926; PubMed Central PMCID: PMCPMC8303422.

26. Xiao Y, Lidsky PV, Shirogane Y, Aviner R, Wu CT, Li W, et al. A defective viral genome strategy elicits broad protective immunity against respiratory viruses. Cell. 2021;184(25):6037–51.e14. Epub 2021/12/02. doi: 10.1016/j.cell.2021.11.023. PubMed PMID: 34852237; PubMed Central PMCID: PMCPMC8598942.

27. Wong CH, Ngan CY, Goldfeder RL, Idol J, Kuhlberg C, Maurya R, et al. Reduced subgenomic RNA expression is a molecular indicator of asymptomatic SARS-CoV-2 infection. Commun Med (Lond). 2021;1:33. Epub 2022/05/24. doi: 10.1038/s43856-021-00034-y. PubMed PMID: 35602196; PubMed Central PMCID: PMCPMC9053197.

28. Gribble J, Stevens LJ, Agostini ML, Anderson-Daniels J, Chappell JD, Lu X, et al. The coronavirus proofreading exoribonuclease mediates extensive viral recombination. PLoS Pathog. 2021;17(1):e1009226. Epub 2021/01/20. doi: 10.1371/journal.ppat.1009226. PubMed PMID: 33465137; PubMed Central PMCID: PMCPMC7846108.

29. Ziegler CGK, Miao VN, Owings AH, Navia AW, Tang Y, Bromley JD, et al. Impaired local intrinsic immunity to SARS-CoV-2 infection in severe COVID-19. Cell. 2021;184(18):4713–33 e22. Epub 2021/08/06. doi: 10.1016/j.cell.2021.07.023. PubMed PMID: 34352228; PubMed Central PMCID: PMCPMC8299217.

30. Ravindra NG, Alfajaro MM, Gasque V, Huston NC, Wan H, Szigeti-Buck K, et al. Single-cell longitudinal analysis of SARS-CoV-2 infection in human airway epithelium identifies target cells, alterations in gene expression, and cell state changes. PLoS Biol. 2021;19(3):e3001143. Epub 20210317. doi: 10.1371/journal.pbio.3001143. PubMed PMID: 33730024; PubMed Central PMCID: PMCPMC8007021.

31. Bouhaddou M, Reuschl AK, Polacco BJ, Thorne LG, Ummadi MR, Ye C, et al. SARS-CoV-2 variants evolve convergent strategies to remodel the host response. Cell. 2023;186(21):4597–614 e26. Epub 20230921. doi: 10.1016/j.cell.2023.08.026. PubMed PMID: 37738970; PubMed Central PMCID: PMCPMC10604369.

32. Syed AM, Taha TY, Tabata T, Chen IP, Ciling A, Khalid MM, et al. Rapid assessment of SARS-CoV-2-evolved variants using virus-like particles. Science. 2021;374(6575):1626–32. Epub 20211104. doi: 10.1126/science.abl6184. PubMed PMID: 34735219; PubMed Central PMCID: PMCPMC9005165.

33. Torii S, Ono C, Suzuki R, Morioka Y, Anzai I, Fauzyah Y, et al. Establishment of a reverse genetics system for SARS-CoV-2 using circular polymerase extension reaction. Cell Rep. 2021;35(3):109014. Epub 2021/04/12. doi: 10.1016/j.celrep.2021.109014. PubMed PMID: 33838744; PubMed Central PMCID: PMCPMC8015404.

34. Pechous RD, Malaviarachchi PA, Banerjee SK, Byrum SD, Alkam DH, Ghaffarieh A, et al. An ex vivo human precision-cut lung slice platform provides insight into SARS-CoV-2 pathogenesis and antiviral drug efficacy. J Virol. 2024;98(7):e0079424. Epub 20240628. doi: 10.1128/jvi.00794-24. PubMed PMID: 38940558; PubMed Central PMCID: PMCPMC11265413.

35. Robinson JT, Thorvaldsdottir H, Winckler W, Guttman M, Lander ES, Getz G, Mesirov JP. Integrative genomics viewer. Nat Biotechnol. 2011;29(1):24–6. doi: 10.1038/nbt.1754. PubMed PMID: 21221095; PubMed Central PMCID: PMCPMC3346182.

36. Offersgaard A, Duarte Hernandez CR, Pihl AF, Costa R, Venkatesan NP, Lin X, et al. SARS-CoV-2 Production in a Scalable High Cell Density Bioreactor. Vaccines (Basel). 2021;9(7). Epub 20210629. doi: 10.3390/vaccines9070706. PubMed PMID: 34209694; PubMed Central PMCID: PMCPMC8310283.

37. Gao Y, Yan L, Huang Y, Liu F, Zhao Y, Cao L, et al. Structure of the RNA-dependent RNA polymerase from COVID-19 virus. Science. 2020;368(6492):779–82. Epub 20200410. doi: 10.1126/science.abb7498. PubMed PMID: 32277040; PubMed Central PMCID: PMCPMC7164392.

38. Luo Y, Yu F, Zhou M, Liu Y, Xia B, Zhang X, et al. Engineering a Reliable and Convenient SARS-CoV-2 Replicon System for Analysis of Viral RNA Synthesis and Screening of Antiviral Inhibitors. mBio. 2021;12(1). Epub 20210119. doi: 10.1128/mBio.02754-20. PubMed PMID: 33468688; PubMed Central PMCID: PMCPMC7845634.

39. Gu L, Wang ZJ, Zhang XR, Liu Y, Zhao M, Jiang SZ, et al. Targeting the liquid-liquid phase separation of nucleocapsid broadly inhibits the replication of SARS-CoV-2 strains. Biochem Biophys Res Commun. 2025;756:151594. Epub 20250306. doi: 10.1016/j.bbrc.2025.151594. PubMed PMID: 40086356.

40. Zheng Y, Deng J, Han L, Zhuang MW, Xu Y, Zhang J, et al. SARS-CoV-2 NSP5 and N protein counteract the RIG-I signaling pathway by suppressing the formation of stress granules. Signal Transduct Target Ther. 2022;7(1):22. Epub 20220124. doi: 10.1038/s41392-022-00878-3. PubMed PMID: 35075101; PubMed Central PMCID: PMCPMC8785035.

41. Bessa LM, Guseva S, Camacho-Zarco AR, Salvi N, Maurin D, Perez LM, et al. The intrinsically disordered SARS-CoV-2 nucleoprotein in dynamic complex with its viral partner nsp3a. Sci Adv. 2022;8(3):eabm4034. Epub 20220119. doi: 10.1126/sciadv.abm4034. PubMed PMID: 35044811; PubMed Central PMCID: PMCPMC8769549.

42. Cubuk J, Alston JJ, Incicco JJ, Singh S, Stuchell-Brereton MD, Ward MD, et al. The SARS-CoV-2 nucleocapsid protein is dynamic, disordered, and phase separates with RNA. Nat Commun. 2021;12(1):1936. Epub 20210329. doi: 10.1038/s41467-021-21953-3. PubMed PMID: 33782395; PubMed Central PMCID: PMCPMC8007728.

43. Desmyter J, Melnick JL, Rawls WE. Defectiveness of interferon production and of rubella virus interference in a line of African green monkey kidney cells (Vero). J Virol. 1968;2(10):955–61. doi: 10.1128/JVI.2.10.955-961.1968. PubMed PMID: 4302013; PubMed Central PMCID: PMCPMC375423.

44. Ricciardi S, Guarino AM, Giaquinto L, Polishchuk EV, Santoro M, Di Tullio G, et al. The role of NSP6 in the biogenesis of the SARS-CoV-2 replication organelle. Nature. 2022;606(7915):761–8. Epub 20220512. doi: 10.1038/s41586-022-04835-6. PubMed PMID: 35551511; PubMed Central PMCID: PMCPMC7612910.

45. Chaturvedi S, Vasen G, Pablo M, Chen X, Beutler N, Kumar A, et al. Identification of a therapeutic interfering particle-A single-dose SARS-CoV-2 antiviral intervention with a high barrier to resistance. Cell. 2021;184(25):6022–36 e18. Epub 2021/11/29. doi: 10.1016/j.cell.2021.11.004. PubMed PMID: 34838159; PubMed Central PMCID: PMCPMC8577993.

46. Shemesh M, Aktepe TE, Deerain JM, McAuley JL, Audsley MD, David CT, et al. SARS-CoV-2 suppresses IFNbeta production mediated by NSP1, 5, 6, 15, ORF6 and ORF7b but does not suppress the effects of added interferon. PLoS Pathog. 2021;17(8):e1009800. Epub 20210826. doi: 10.1371/journal.ppat.1009800. PubMed PMID: 34437657; PubMed Central PMCID: PMCPMC8389490.

47. Brandherm L, Kobas AM, Klohn M, Bruggemann Y, Pfaender S, Rassow J, Kreimendahl S. Phosphorylation of SARS-CoV-2 Orf9b Regulates Its Targeting to Two Binding Sites in TOM70 and Recruitment of Hsp90. Int J Mol Sci. 2021;22(17). Epub 20210826. doi: 10.3390/ijms22179233. PubMed PMID: 34502139; PubMed Central PMCID: PMCPMC8430508.

48. Ghosh S, Dellibovi-Ragheb TA, Kerviel A, Pak E, Qiu Q, Fisher M, et al. beta-Coronaviruses Use Lysosomes for Egress Instead of the Biosynthetic Secretory Pathway. Cell. 2020;183(6):1520–35 e14. Epub 20201027. doi: 10.1016/j.cell.2020.10.039. PubMed PMID: 33157038; PubMed Central PMCID: PMCPMC7590812.

49. Tilocca B, Soggiu A, Sanguinetti M, Musella V, Britti D, Bonizzi L, et al. Comparative computational analysis of SARS-CoV-2 nucleocapsid protein epitopes in taxonomically related coronaviruses. Microbes Infect. 2020;22(4-5):188–94. Epub 20200414. doi: 10.1016/j.micinf.2020.04.002. PubMed PMID: 32302675; PubMed Central PMCID: PMCPMC7156246.

50. Li Y, Renner DM, Comar CE, Whelan JN, Reyes HM, Cardenas-Diaz FL, et al. SARS-CoV-2 induces double-stranded RNA-mediated innate immune responses in respiratory epithelial-derived cells and cardiomyocytes. Proc Natl Acad Sci U S A. 2021;118(16). doi: 10.1073/pnas.2022643118. PubMed PMID: 33811184; PubMed Central PMCID: PMCPMC8072330.

51. Pryhuber GS, Huyck, H., . URMC_HTC_Whole Lung and Lobe Processing. (2020) [cited 603.3 & 604.5].

52. Pitonzo A CA, Huyck H, Pryhuber GS. HTC_Precision_Cut_Lung_Slices V.2. 2025. [cited 623.2.].

53. Zheng GX, Terry JM, Belgrader P, Ryvkin P, Bent ZW, Wilson R, et al. Massively parallel digital transcriptional profiling of single cells. Nat Commun. 2017;8:14049. Epub 2017/01/17. doi: 10.1038/ncomms14049. PubMed PMID: 28091601; PubMed Central PMCID: PMCPMC5241818 L.M., D.A.M., S.Y.N., M.S.L., P.W.W., C.M.H., R.B., A.W., K.D.N., T.S.M. and B.J.H. are employees of 10x Genomics.

54. Satija R, Farrell JA, Gennert D, Schier AF, Regev A. Spatial reconstruction of single-cell gene expression data. Nat Biotechnol. 2015;33(5):495–502. Epub 2015/04/14. doi: 10.1038/nbt.3192. PubMed PMID: 25867923; PubMed Central PMCID: PMCPMC4430369.

55. Smith T, Heger A, Sudbery I. UMI-tools: modeling sequencing errors in Unique Molecular Identifiers to improve quantification accuracy. Genome Res. 2017;27(3):491–9. Epub 2017/01/20. doi: 10.1101/gr.209601.116. PubMed PMID: 28100584; PubMed Central PMCID: PMCPMC5340976.

56. Langmead B, Salzberg SL. Fast gapped-read alignment with Bowtie 2. Nat Methods. 2012;9(4):357–9. Epub 2012/03/06. doi: 10.1038/nmeth.1923. PubMed PMID: 22388286; PubMed Central PMCID: PMCPMC3322381.

57. di Sant’Agnese PA, De Mesy-Jensen KL. Diagnostic Electron Microscopy on Reembedded (“Popped Off”) Areas of Large Spun* Epoxy Sections. Ultrastructural Pathology. 1984;6(2-3):247–52. doi: 10.3109/01913128409018580.

